# High-throughput phenotyping reveals expansive genetic and structural underpinnings of immune variation

**DOI:** 10.1101/688010

**Authors:** Lucie Abeler-Dörner, Adam G. Laing, Anna Lorenc, Dmitry S. Ushakov, Simon Clare, Anneliese Speak, Maria Duque, Jacqui K. White, Ramiro Ramirez-Solis, Namita Saran, Katherine R. Bull, Belén Morón, Jua Iwasaki, Philippa R. Barton, Susana Caetano, Keng I. Hng, Emma Cambridge, Simon Forman, Tanya L. Crockford, Mark Griffiths, Leanne Kane, Katherine Harcourt, Cordelia Brandt, George Notley, Kolawole O. Babalola, Jonathan Warren, Jeremy C. Mason, Amrutha Meeniga, Natasha A. Karp, David Melvin, Eleanor Cawthorne, Brian Weinrick, Albina Rahim, Sibyl Drissler, Justin Meskas, Alice Yue, Markus Lux, George Song-Zhao, Anna Chan, Carmen Ballesteros Reviriego, Johannes Abeler, Heather Wilson, Agnieszka Przemska-Kosicka, Matthew Edmans, Natasha Strevens, Markus Pasztorek, Terrence F. Meehan, Fiona Powrie, Ryan Brinkman, Gordon Dougan, William Jacobs, Clare Lloyd, Richard J. Cornall, Kevin Maloy, Richard Grencis, Gillian M. Griffiths, David Adams, Adrian C. Hayday

**Affiliations:** Department of Immunobiology, King’s College London, UK; The Francis Crick Institute, London, UK; Wellcome Sanger Institute, Hinxton, UK; MRC Human Immunology Unit, University of Oxford, UK; Dunn School of Pathology, University of Oxford, UK; Imperial College London, UK; University of Cambridge, UK; University of Manchester, UK; European Bioinformatics Institute, Hinxton, UK; Albert Einstein College of Medicine, New York, US; University of British Columbia, Vancouver, Canada; Department of Economics, University of Oxford, UK; The Kennedy Institute of Rheumatology, University of Oxford, UK

## Abstract

By developing a high-density murine immunophenotyping platform compatible with high-throughput genetic screening, we have established profound contributions of genetics and structure to immune variation. Specifically, high-throughput phenotyping of 530 knockout mouse lines identified 140 monogenic “hits” (>25%), most of which had never hitherto been implicated in immunology. Furthermore, they were conspicuously enriched in genes for which humans show poor tolerance to loss-of-function. The immunophenotyping platform also exposed dense correlation networks linking immune parameters with one another and with specific physiologic traits. By limiting the freedom of individual immune parameters, such linkages impose genetically regulated “immunological structures”, whose integrity was found to be associated with immunocompetence. Hence, our findings provide an expanded genetic resource and structural perspective for understanding and monitoring immune variation in health and disease.

---

Because immune function is increasingly implicated in almost every arena of pathophysiology, there is growing demand for more insight into the basis of inter-individual immune variation and for incisive ways to measure it at steady-state and in response to challenges and treatments. Reflecting this are many highly informative studies describing human immune system dynamics^1–4^, and investigations of the factors contributing to it^5–8^. Thus, SNP-based and deep sequencing-based Genome-Wide Association Studies (GWAS) and Twin-studies have made compelling associations of defined genetic loci with autoimmunity and/or immunodeficiency^9–14^. However, there has been much discussion of the difficulties of directly linking specific genes and/or genetic variants to discrete immunophenotypes^15^. Conversely, concrete links between specific genes and immune function have emerged from the analysis of Mendelian traits, an approach that is expanding through genome sequencing of many “rare diseases”^16, 17^. Nonetheless, this approach can be limited by the infrequency of diseases, uncertain clinical annotation, and limitations on the phenotypic breadth that can be characterized.

At the same time, it has become clear that sex, age, and environmental factors, including diet and the microbiome, make major contributions to human immune variation^5, 6, 8^, but assessing their full impacts is limited by appropriate constraints on interventions, and by the genetic diversity of human populations. In sum, there are so many diverse immunoregulatory factors that it becomes difficult to understand to what degree and by which means any single factor impacts upon the immune system.

In this regard, animal model studies offer unique opportunities, despite the limitations on their extrapolation to human pathophysiology. Specifically, use of an inbred strain limits genetic variation; co-housing reduces microbiome and dietary variation; and the use of age-matched animals limits physiologic variation. By this means, their study can establish a template for the nature and sources of variation in the “baseline immune system”, thereby directly informing myriad investigations of rodent immunology and usefully guiding the design and interpretation of human immune system studies.

Added to this, a high-throughput genetic screen of co-housed, age-matched mice could provide clear insight into the fraction of genes whose loss-of-function perturbs one or more aspects of the baseline immune system, and the nature and functional consequences of those perturbations. Such insight could be expected to reveal novel pathways and forms of immunoregulation relevant to myriad discrete endophenotypes (e.g. prevalence or activity of a defined immune cell type), which can be extremely useful in linking genetic variants to disease^15^. By better depicting the basis and nature of immune variation, such insight could also guide the development of more incisive, practical approaches to immune monitoring. Furthermore, because many diverse phenotypes can be measured in mice, the opportunity exists to relate immune variation to discrete physiologic traits.

To achieve these goals at scale, we have developed a robust, routinely applicable, high-density, high-throughput Infection and Immunity Immunophenotyping [“3i”] platform (www.immunophenotype.org) that has in under 5 years permitted the analysis of the baseline immune system and response to challenge in 530 isogenic, aged-matched, co-housed mutant mouse strains. Additionally, by integrating 3i into the International Mouse Phenotyping Consortium (IMPC) pipeline (www.mousephenotype.org), immune variation could be related to many measures of general physiology.

The expansive outputs of this experimental approach (>1million data-points) have provided several discrete insights and data-rich resources that collectively offer a revised frame-of-reference for viewing immune variation. Specifically, we found that variation in the baseline immune system was contributed to differentially by discrete immune subsets, many of which showed overt sexual dimorphism. Furthermore, variation in specific baseline immune parameters was commonly correlated, positively or negatively, with variation in other immune parameters. While some such correlations were expected, many were not. Additionally, variation in specific immune subsets also correlated with variation in specific, non-immune, physiological traits such as HDL-cholesterol or fructose. By inevitably limiting the freedom-of-movement of any single immune parameter, such correlations must ordinarily impose structural constraints on immune variation.

Of the 530 genes screened, the baseline immunophenotype and/or responses to challenge were affected by mutations in 140 (>25%) genes [a.k.a. “hits”]. Of note, 57% of hits were not hitherto implicated in immunobiology, and yet they were strikingly enriched in genes for which humans show little tolerance of loss-of-function. Hence, 3i has greatly expanded the number and diversity of genes implicated in immune system biology, with effects ranging from impacts on single parameters to broad immune dysregulation. Furthermore, genes whose mutation affected several immune subsets could be classified into those that substantially preserved the correlations between immune subsets *versus* those that largely disrupted the resulting structures. Interestingly, genes that affected immune and non-immune traits caused significantly greater structural disruption than did genes that only affected immune traits. Moreover, greater disruption was also caused by those genes affecting challenge responses as well as the baseline immune system, thereby associating the integrity of immune structures with host defense. Hence, immunological structure can be an important consideration in the design of immune monitoring.

## Results

### Immunophenotyping of mutant mice at scale

The goal of the International Mouse Phenotyping Consortium (IMPC) (www.mousephenotype.org) has been to generate and provide open-access phenotyping data for mice with targeted disruptions in each of ∼18,000 annotated protein-coding genes, obtained using embryonic stem cells generated by the International Knockout Mouse Consortium^18^ and, more recently, using CRISPR technology. As a major contributor to IMPC, the Wellcome Trust Sanger Institute (WTSI) generated approximately three new mutant lines per week. Given this scale, immunological assays performed by IMPC were limited to peripheral blood lymphocytometry and responses to *Salmonella* and *Citrobacter* infection^19^ (**Fig 1A; Fig S1A**), inevitably failing to score many genes contributing to immune variation. Therefore, a high-density infection and immunity immunophenotyping platform (3i) was developed (**Fig 1A**) to be compatible with the IMPC high-throughput screen (HTS). At one and the same time, this offered the potential to greatly expand the number and diversity of genes implicated in immunobiology, and to provide a better estimate of the fraction and types of genes that may underpin immune variation. Moreover, by integration into the IMPC, 3i could relate immunophenotypes to many non-immunological traits.

**Figure 1:**
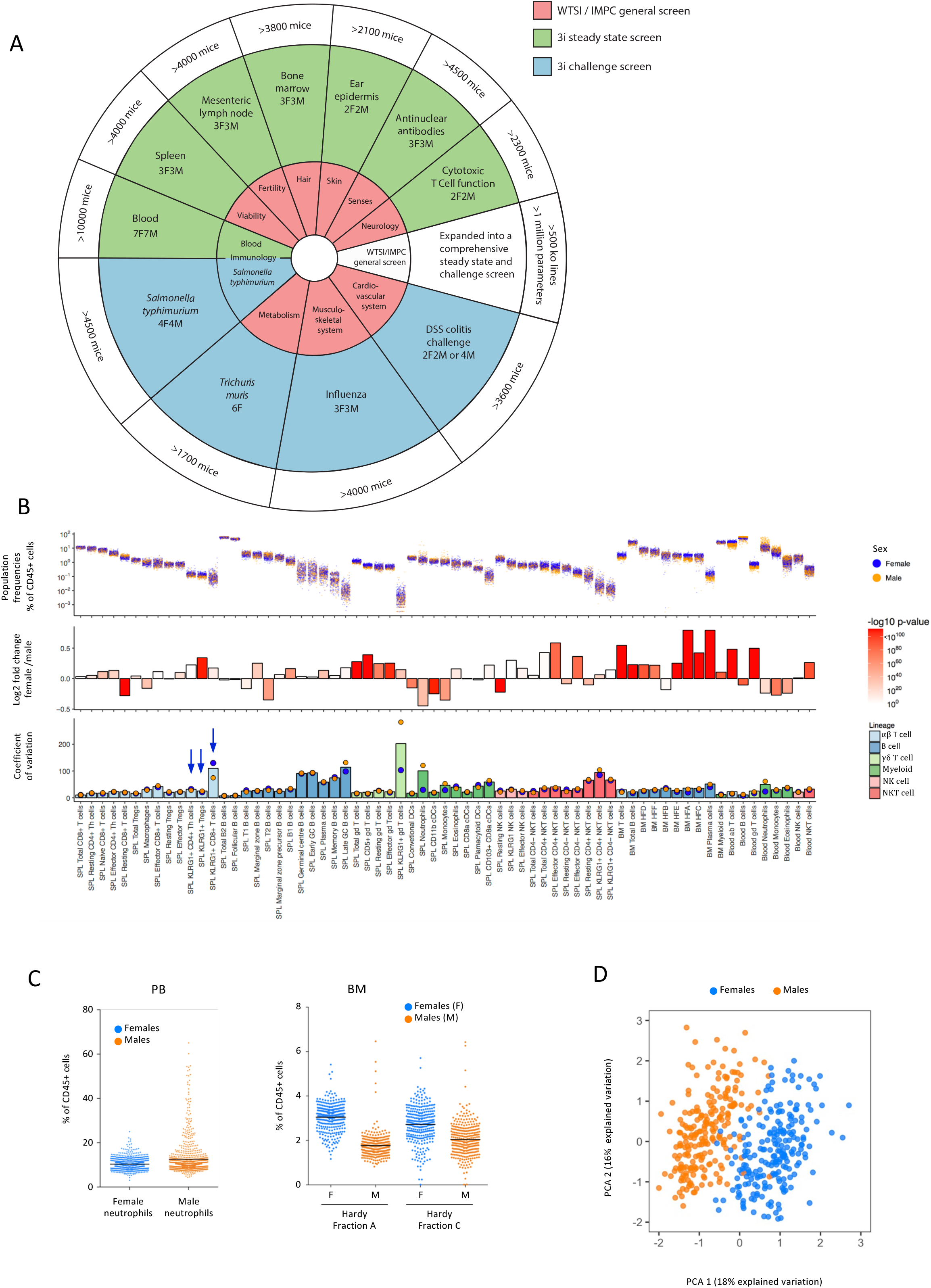
Variation in immune cell subset composition with sex as a contributory driver. A. Overview of the tests performed by WTSI as part of IMPC (inner circle) and by 3i and WTSI as part of 3i (outer circle). B. Sexual dimorphism is wide-spread in the immune system: Population sizes as % of CD45 cells (upper panel); sexual dimorphism of mean values of population sizes as log2 fold change (middle panel, female/male); and coefficient of variation (lower panel) of immune populations in SPL, BM and PB from 16-week old male and female wt C57BL/6N mice (n>500). Blue arrows denote cell subsets mentioned in text. C. Left: Neutrophils in peripheral blood of 16-week old wt C57BL/6N mice (n>900 per sex) show sexual dimorphism in population frequency (bars represent means) and in variance. Right: Hardy fractions A and C in the bone marrow of 16-week old WT C57BL/6n mice (n>280 per sex) show sexual dimorphism in population frequencies. D. Data on 60 immune cell subset frequencies from 451 wt mice (one dot each) was sufficient to segregate sexes with 99% accuracy. PCA of cell type frequencies from four tissues (SPL, MLN, BM, PB); first two PCA axes explain 34% of total variation; colour denotes sex.

At homeostasis, the baseline immune system is simultaneously poised to respond to infectious or toxic challenges and regulated to limit immunopathology. Hence, inter-individual variation in this state is likely manifest in differential immunocompetence and susceptibilities to autoimmune diseases. To capture this, a high-content flow cytometry analysis of lymphoid and myeloid cells and their activation states in spleen (SPL), mesenteric lymph nodes (MLN), bone marrow (BM) and peripheral blood (PB) was performed at steady-state (**Fig 1A;** panels in **Table S1**; populations quantitated in **Table S2**; illustrative gating strategy for MLN T, NKT, and NK cell subsets shown in **Fig S1B**; gating strategies for all other panels shown in **Fig S4**). To sample the immune system in an extra-lymphoid tissue, quantitative object-based imaging was applied to intra-epidermal lymphoid and myeloid cells *in situ* (**Fig S1C**). Anti-nuclear antibodies (ANA) were quantitated since they commonly reflect impaired immunological tolerance (**Fig S1D**), while effector potential was gauged by measuring cytolysis by SPL CD8 T cells.

To be compatible with the IMPC, immunological assays could not divert tissue from, or operationally impinge upon the basic phenotyping programme. Thus, all observational assays requiring sacrifice were conducted at the prescribed termination-point for IMPC assays of 16 weeks. In parallel, mice were assayed for responses to infection by a parasite (*Trichuris muris*), a virus (influenza), and a bacterium (*Salmonella typhimurium*), and to sodium dextran sulphate (DSS) that causes gut epithelial erosion and microbial translocation (**Fig 1A**; **Fig S1E**). For each component of 3i, experimental standard-operating procedures were established and stringently quality-controlled; instruments were well calibrated; and data reproducibility monitored longitudinally, with automated analysis accounting for any temporal variation (**Fig S1F**; see **Materials and Methods**). Additionally, and to be compatible with the time-and- budget constraints of an HTS, we chose minimum numbers of data-points required to establish significance following application of bespoke statistical analyses (**Fig 1A;** **Table S3**).

This study reports on the first five-year phase in which 3i phenotyped 530 mutant mouse strains (**Table S4**; see also www.immunophenotype.org): most were knockouts/nulls or severe hypomorphs of protein-coding genes, whereas 1.8% were lncRNA/miRNA mutants. For 30% of the genes, heterozygotes were screened because homozygotes were embryonically lethal or sub-viable. Of the genes selected, ∼30% were chosen because they were linked to a disease, e.g. by GWAS, whereas to maximize discovery, the majority were mostly little-studied genes chosen either randomly or guided by investigators’ interests.

Overall, >1 million data-points were collected from 7 distinct steady-state assay systems applied to 2,100 −10,000 mice (**Fig 1A**), while several thousand additional mice were subjected to challenges. Of note, the minimization of technical variation; fastidious control for batch variation; optimization of data collection and analysis^20^; and innovative data management across heterologous platforms permitted 3i to make rigorous assessments of naturally-arising variation in the baseline immune system of many hundreds of genetically identical, age-matched, co-located, adult wild-type (wt) “control” C57.BL/6N mice, thereby creating a precise backdrop for the analysis of mutants.

### Immunophenotypic variation among controls

Most steady-state immune cell subsets in adult C57.BL/6N controls showed low coefficients of variation (CV), although these were further reduced by dynamic automated gating of flow cytometry data, particularly for numerically small cell subsets whose reproducible quantitation can be challenging (**Fig S1G**). Hence, automated gating was adopted screen-wide to obtain population sizes and CVs (**Figs 1B;** see **Materials and Methods**)^20^. This revealed greater variation for some specific cell types, including germinal centre (GC) B cells and various αβ and γδ T cell subsets expressing the activation marker KLRG1 (**Fig 1B**, bottom-most panel). Activation-driven variation of adaptive immune subsets was anticipated since non-heritable, antigen receptor gene rearrangements dictate different responses of syngeneic individuals to shared environments. Nonetheless, the effects were highly selective, as evidenced by low CVs of KLRG1^+^ CD4 T helper cells and of KLRG1^+^ Treg cells compared to a comparably-sized subset of KLRG1^+^ CD8 T cells (**Fig 1B**, compare top and bottom-most panels for subsets denoted with blue arrows).

Most innate immune subsets showed relatively low CVs, although SPL and PB neutrophils were exceptions, particularly in male mice. Indeed, sex was a consistent source of variation for ∼50% of PB, SPL, BM, and MLN cell subsets, reflected in either significantly different variance for female (F) and male (M) mice, as illustrated for PB neutrophils (**Fig 1C**; left panel) and displayed broadly in **Fig 1B** (bottom panel - CV^F^, blue circles; CV^M^, orange circles), and/or in sexually dimorphic mean values, as illustrated for BM B cell progenitors (**Fig 1C**; right panel), and displayed broadly in **Fig 1B** (middle panel; mean^F^/mean^M^ log2-transformed). Indeed, the impact of sex was so profound that principal component analysis (PCA) of sixty aggregated SPL, MLN, BM, and PBL flow cytometry parameters was sufficient to sex-segregate 451 mice with >99% accuracy (**Fig 1D**).

Cell counts and several other properties of Vγ5^+^ dendritic epidermal T cells (DETC) and Langerhans Cells (LC) were also significantly sexually dimorphic, as were ANA outputs, evoking the frequent gender imbalance of human autoimmunity (**Fig S1H)**^21^. Conversely, DSS outputs did not segregate significantly by sex, possibly germane to gender-neutral incidences of inflammatory bowel disease (**Fig S1H**)^22^. In sum, the baseline immune system of adult C57.BL/6 mice, the most frequently used strain in immunological investigations, showed widespread but highly selective sexual dimorphism. As a practical response to this, all statistical tests of 3i data accounted for sex (see **Materials and Methods**).

### Significant correlations of discrete immune parameters

Although the immune system clearly comprises a dynamic, multi-component ecosystem, the inter-connectedness of its constituent cell populations is poorly understood. In this regard, the 3i analysis of >650 age-matched, co-located, genetically identical control mice identified significant positive (red) and negative (blue) correlations, as illustrated for 46 steady-state SPL parameters in male and female mice (**Fig 2A**; **Fig S2A**): note, we excluded contingent relationships reflecting nested or paired technical correlates (see **Fig S1B; Fig S4)**.

**Figure 2:**
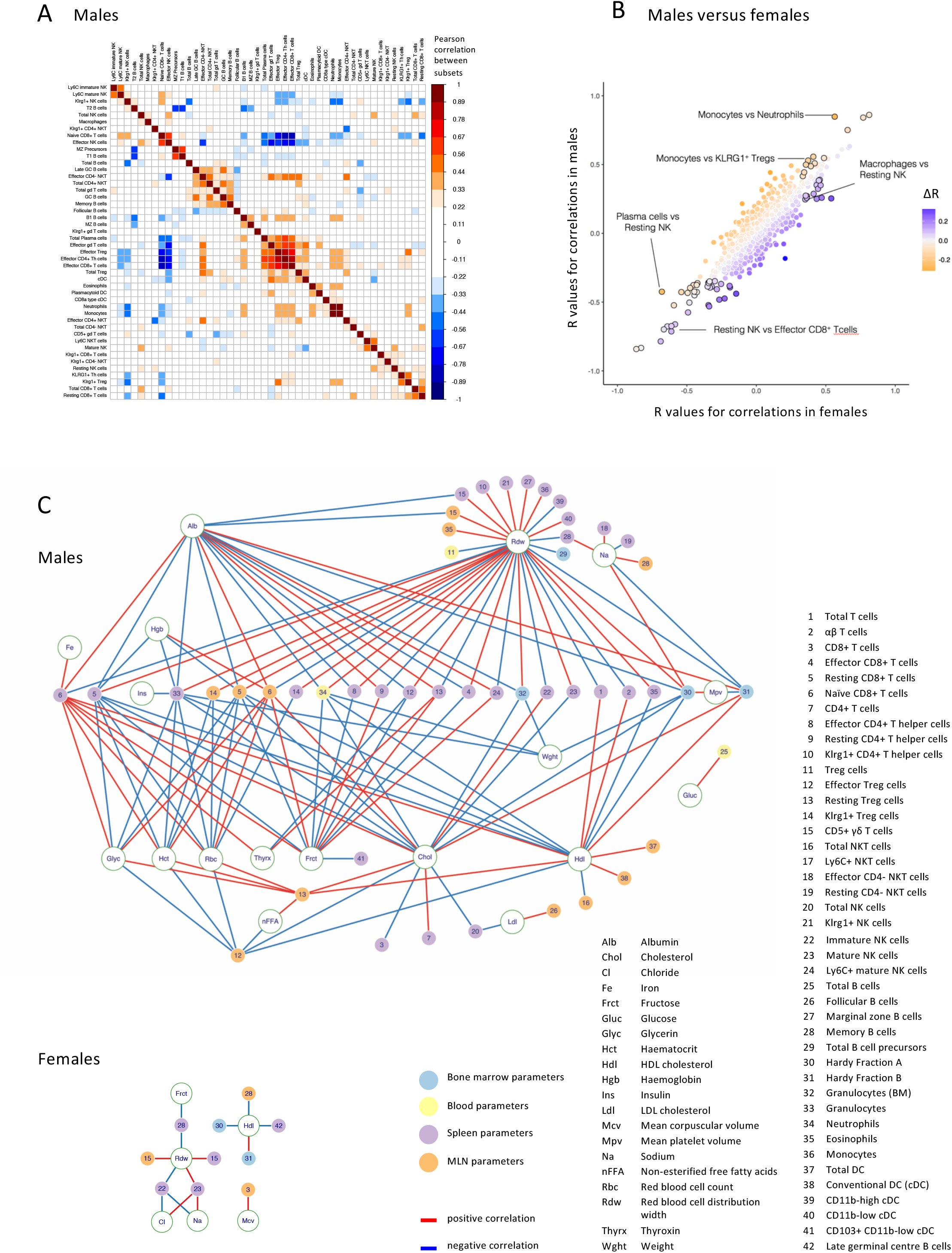
Correlations between immune and non-immune parameters reveal a sex-specific network of interactions. A Several SPL cell subsets correlate with each other. Heat map represents Pearson correlations of 46 splenic immune cell subsets with each other in wt males (n>230) as determined by flow cytometry. Dark red fields denote strong positive, dark blue fields strong negative correlations between frequencies of spleen immune cell subsets. B Specific correlations differ between males and females. Colour denotes ΔR, the difference between the correlation coefficient R for SPL subsets in male and female wt mice. Parameter pairs that are significantly sexually dimorphic have black circumference (see **Materials and Methods**). Correlation coefficients for male and female mice were derived from data depicted in Fig 2A and Fig S2A, respectively. C The correlation network for male immune and nonimmune parameters is more pronounced than for females. Depicted are correlations with a Pearson R-value >0.33 and p<0.001 between PBL, SPL, MLN, BM with haematological, clinical blood chemistry, and additional parameters (see **Fig S2B**). Circle colours denote organ assayed; red lines denote positive correlations; blue lines, negative correlations (n>180 per sex).

As illustrated by the male and female SPL data-sets (**Fig 2A**, **Fig S2A**), correlations fell mostly into three classes, of which the first related to activation state. Thus, a “lymphoid activation-cluster” embraced effector CD4^+^, CD8^+^, NK and *γδ* T cells; Tregs; conventional DC; and plasma cells (**Fig 2A**, **Fig S2A**; central red core). Some such relationships were anticipated: e.g. effector Th correlated with total Treg, and even better with effector Treg. Conversely, other correlations were unanticipated; e.g. effector CD8^+^ T cells correlated positively with plasma cells (**Fig 2A**, 27 down, 23 across) but negatively with effector NK cells (**Fig 2A**, 27 down, 9 across). Such relationships have implications for the design and monitoring of vaccines aimed at eliciting discrete effector responses.

A second class of correlations related to immune homeostasis. For example, higher NK cell representation reflected increases in mature NK cells (**Fig 2A**, 5 down, 9 across), whereas higher CD8^+^ T cell representation reflected increases in resting but not effector CD8^+^ T cells (**Fig 2A**, 45 down *versus* 46 across or 27 across). Finally, a third class of correlations comprised relationships not easily attributable to current knowledge: e.g. significant positive relationships of monocytes with B1 B cells, and of memory B cells with *γδ* T cells (**Fig 2A**, 34 down, 20 across; 18 down, 16 across). Exploring the basis of such relationships offers new avenues for immunological investigation.

Highlighting their robustness, many of the identified relationships were largely comparable in male and female mice (**Fig 2A**, **Fig S2A**; **colorless** circles along the 45° axis in **Fig 2B**). Nonetheless, some parameter pairings were sexually dimorphic, with R values and/or regression slopes being stronger in females (e.g. macrophages *versus* resting NK) or *vice versa* (e.g. monocytes *versus* neutrophils) (**Fig 2B**, purple and orange circles, respectively). Whereas the spleen matrices shown are intra-organ, there were likewise many inter-organ correlations (see below), collectively revealing the baseline immune system of adult C57.BL/6 mice to be underpinned by dense, sexually dimorphic networks of >1000 correlations. Possibly these reflect robust intercellular circuitry, as was recently illustrated for macrophages and fibroblasts^23^.

### Integrating immunology with physiology

By coupling 3i to the IMPC, all animals in the observational screen were subject to measures of general physiology (see **Figs 1A; S1A**). As is well established, many such measures exhibited positive or negative correlations with each other, which by limiting the freedom-of-movement of individual parameters, imposed dimorphic phenotypic structures for female and male mice that reflect sex-specific physiologies (**Fig S2B**). Of note, the correlation network appeared much less dense in females, possibly because its full elucidation was prevented by the additional variation arising from sex-specific components such as oestrous that cycles every 4-5 days^24^. This notwithstanding, some core relationships were clearly conserved in females and males, e.g. positive correlations spanning cholesterol (Chol), HDL-cholesterol (Hdl), insulin (Ins), and body-weight (Wght), and the relationship of total protein (Tp) to fructosamine (Fruct), which reflects blood protein glycation. Likewise, glucose (Gluc) correlated negatively with chloride (Cl) and sodium (Na). From a practical standpoint, prominent correlations did not simply reflect high dynamic ranges of specific measurements, since Na was among the least variable.

The more consistent variation of non-immunological parameters in males *versus* females allowed us to identify many highly significant correlations between those non-immunological parameters and specific immunological parameters, with Chol, Hdl, and Na again being prominent (**Fig 2C**). Although immunoregulatory roles for metabolic products and processes are well documented^25, 26^, the correlations identified here were conspicuously selective: thus, there was no overt immunophenotypic correlation with triglycerides, creatinine, or calcium. Interestingly, many correlations were with red blood cell distribution width (Rdw), a common marker of anaemia^27^, frequently associated with chronic immune activation^28, 29^, and recently shown to predict all-cause mortality in humans^30^.

Among immune components, CD8^+^ T cells displayed 40 statistically significant correlations in male mice, compared to 26 for Tregs; 20 for granulocytes, including blood neutrophils; 10 for NK cells; 8 for CD4^+^ T cells (excluding Tregs); and 4 for γδ T cells. These correlations offer insights into physiologic and environmental factors, such as hormones and diet, that may regulate specific immune subsets, and may additionally provide practical surrogate measures of immune status.

Clearly, significant correlations of immune parameters with one another and with discrete non-immunological traits will limit the freedom-to-vary of any single immune parameter, thereby imposing an immunological structure, as shown for males (**Fig 2C**). Because of the sparse correlation network of non-immunological parameters in female mice, a comparable immunological structure was harder to discern (**Fig S2A**; lower panel). Nonetheless, an outline structure emerged, within which Hdl, Rdw and Na were again prominent (**Fig 2C**). Indeed, immunological structure may reflect an important balance of immune and non-immunological components that delivers immunocompetence while limiting immunopathology.

### Many diverse genes affect the immunophenotype

The establishment of robust ranges for myriad immune parameters across age-matched, co-located, wt control mice, coupled with the application of bespoke statistical tests (see **Figs 1A**; **S1F**; **Table S3**; **Materials and Methods**) facilitated high-throughput screening of 530 mutant lines for “phenodeviants” (“hits”) that exerted monogenic dysregulation of the immunophenotype (**Fig 3**). For steady-state immune parameters, phenodeviants were called when ≥60% of samples from a strain located outside the 95^th^ percentiles of the wt distribution for that parameter. To further limit any residual potential for batch effects, each mutant strain was assayed on ≥2 separate occasions, with hits called in a supervised manner (i.e. no data-points were discarded but those possibly attributable to a batch effect would not be scored). False positive rates (FPRs) for each parameter were estimated by randomly sampling sets of wt controls in accordance with the real work flow so as to mimic 10,000 different mutant strains, and then assessing their phenodeviance (**Fig S3A-D**). The resulting FPRs were far below the observed hit-rates that collectively identified 140 phenodeviants, an overall hit-rate of >25% (**Fig S3**). Approximately ∼20% of the hits were heterozygotes, that collectively accounted for ∼30% of the strains analysed; hence, the hit-rate for heterozygotes was ∼18%.

**Figure 3:**
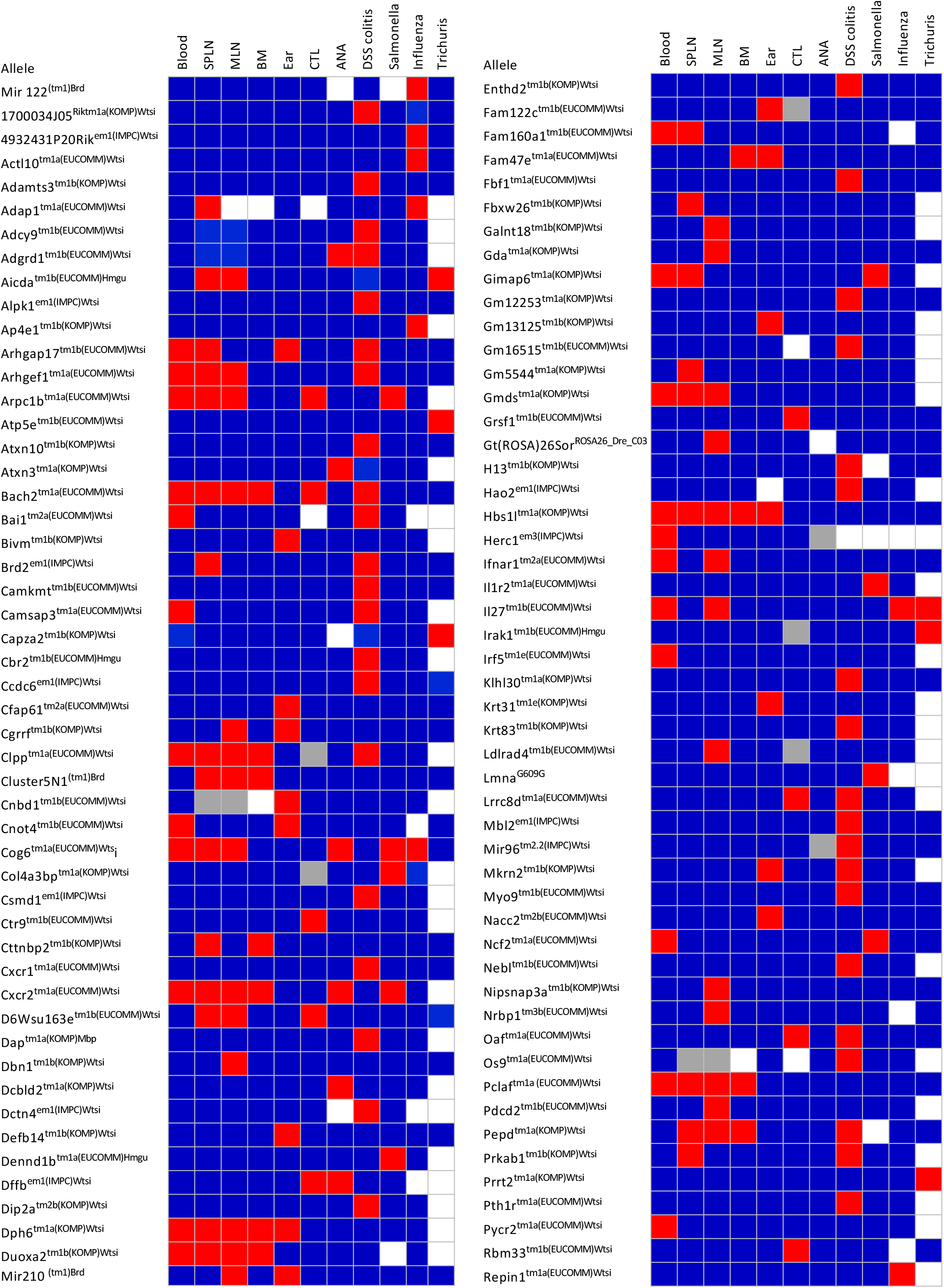
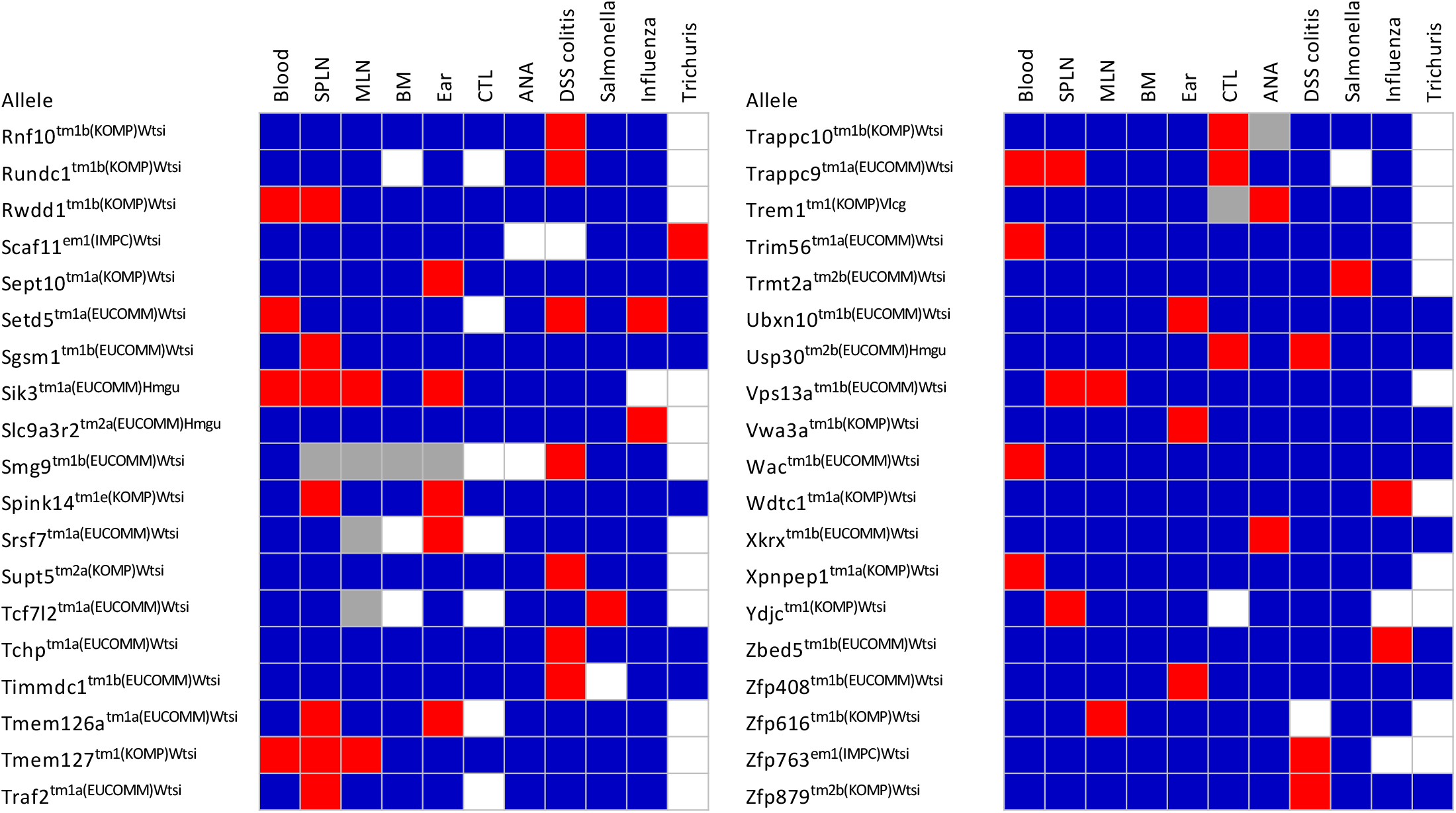
140 out of 530 lines perturb the immunophenotype. Red, significantly different from wt; blue, not significantly different from wt; white, not performed; grey, insufficient data to score as phenodeviant.

Overall hit-calling by 3i was conservative (**Table S3**). Also, strictly sexually dimorphic phenodeviants would not often have scored because with the exceptions of PB cytometry and ANA, the number of mutant mice examined was too small. Nonetheless, sexually dimorphic trends were apparent, for example, in the TCR*αβ*^+^ CD8 T cell percentages in *Arpc1b*^-/-^ mice; in TCR*αβ*^+^ Treg percentages in *Setd5*^-/-^ mice; and in the TCR*αβ*^+^ CD4^+^ CD44^+^ CD62L^-^ effector T cell percentages which are much lower in male *Il27^-/-^* mice (see www.immunophenotype.org).

Because most phenodeviants scored in only one or few assays (**Fig 3**), 3i might also have identified more hits if more immunological assays had been performed. For example, no direct assessments of immunological memory were made, and the steady-state gut could not be analysed because of interference with IMPC phenotyping. Thus, a 25% hit-rate may under-estimate the potential contribution of genetics to immune variation. Moreover, no thorough analysis of the likely genetic impact of non-coding regions, e.g. enhancers, was made.

Conversely, the hit-rate might have been slightly inflated because ∼30% of genes were chosen based on GWAS and/or other implications in immunobiology. Arguing against this however, the hit-rate for little-studied genes was 23% (103/444), closely comparable to all genes; indeed, 3i identified monogenic impacts on the C57.BL/6 immune system of 80 genes never hitherto implicated in immunoregulation. Phenodeviants embodied diverse functions and potentials, collectively ranging from effects on single endophenotypes to very broad perturbations of the immunophenotype. Illustrating this are four case-studies, briefly considered below (**Figs 4, 5**). Note, however, that detailed data on all phenodeviants are provided at www.immunophenotype.org).

### Different classes of genetically regulated immune variation

*Nacc2* (Nucleus accumbens-associated protein 2) is a barely-studied gene^31^ whose highly specific immunological impact was on V*γ*5^+^ DETC numbers (**Fig 4A,B**), a phenotype hitherto observed in C57.BL/6 mice only if they were mutant for V*γ*5 or for *Skint* genes that encode poorly understood, epithelial, DETC-selecting elements^32, 33^. Thus, the 3i screen identified a novel immune regulator that may offer insight into the mechanism by which *Skint* gene-products select and/or maintain a signature tissue-resident T cell compartment.

**Figure 4:**
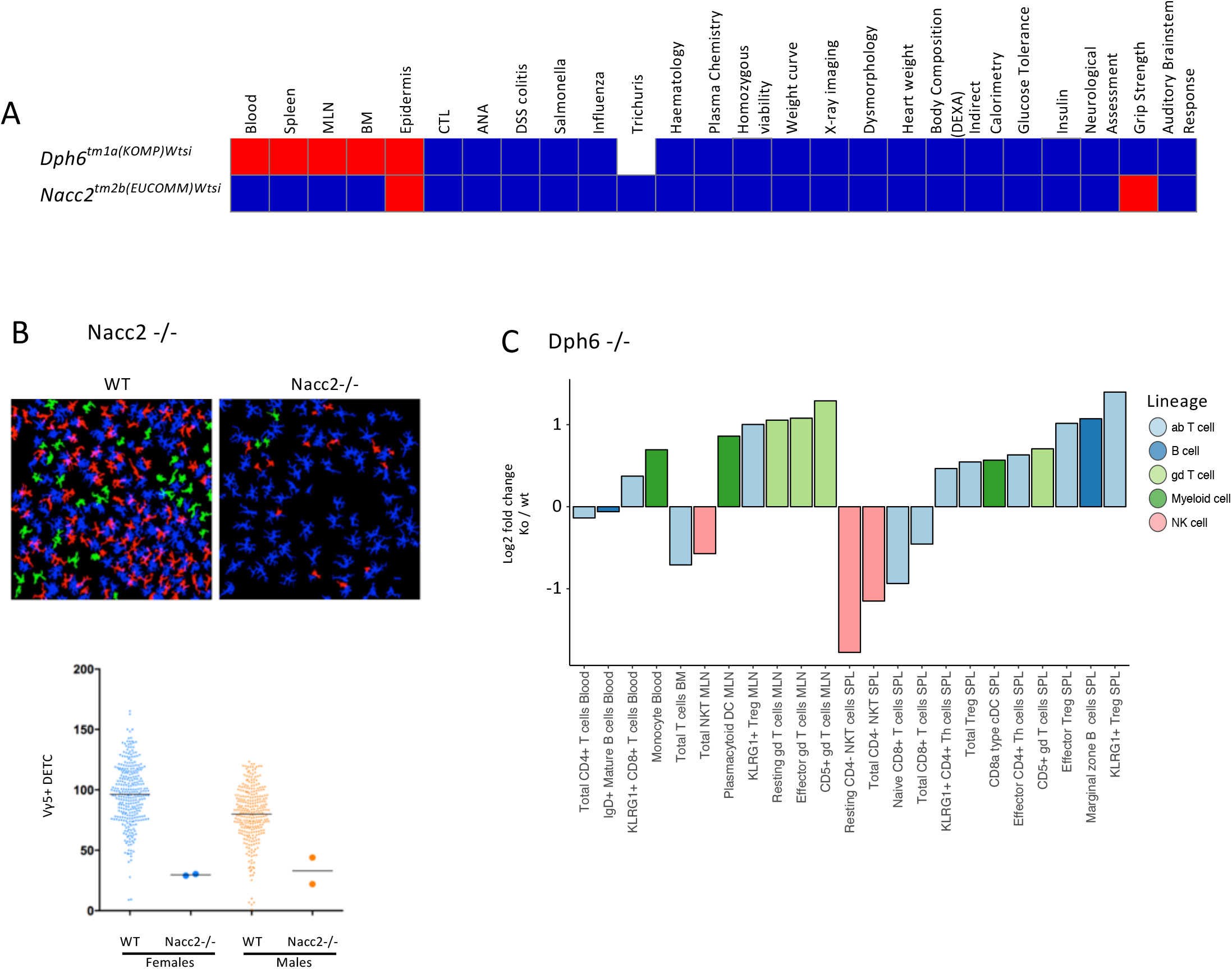
Examples of genes with very specific impacts on the immune system. A. Overview of two genes that display specific immunophenotypes. Columns on the heat map represent different tests: red, significantly different from wt; blue, not significantly different from wt; white, not performed. B. *Nacc2*^-/-^ mice display a specific loss of canonical Vγ5 DETC in ear epidermis. The image represents a z-projection of cell outlines produced by image processing in Definiens Developer XD, which were used for quantitative object-based image analysis: blue, Langerhans cells (LC); red, Vγ5^+^ DETC contacting LC; green: Vγ5^+^ DETC not contacting LC. Bottom: cumulative data for *Nacc2*^-/-^ mice (n=4) versus sex-matched wt controls (n>350 per sex). C. *Dph6^-/-^* mice present with a broad set of hits limited to the immune system. Fold change in immune cell subset proportions between *Dph6^-/-^* and wt mice across multiple tissues (*Dph6^-/-^*, n=6 for SPL, MLN, BM and n=14 for PBL; wt, n>500).

*Dph6^-/-^* mice displayed an immune-specific but broad phenotype, with several innate and adaptive cell lineages showing significant differences from wt (**Fig 4A,C**). Interestingly, *Dph6* is ubiquitously expressed, encoding one of seven enzymes required for synthesizing diphthamide, a modified histidine residue incorporated into eukaryotic elongation factor 2^34^. However, whereas *Dph1, 3* or *4* deficiencies are embryonically lethal^35, 36^, the 3i phenotype suggested that *Dph6* contributes to a previously unreported aspect of immune cell-specific protein translation that is differentially critical for discrete leukocyte subsets. This is germane to growing evidence that cell type-specific translation can be a profound immunoregulator^37^.

In the third case, the broad phenotypic perturbation of several physiologic traits in *Duoxa2*^-/-^ mice (**Fig 5A**) was unsurprising given that Duoxa2 contributes critically to iodine utilisation in thyroid hormone synthesis^38^. By contrast, because, the gene was not hitherto implicated in immunobiology, the significant expansions of specific immune cell subsets, particularly CD4^+^ *αβ* T cells, neutrophils and eosinophils, and the decreased representation of blood Ly6C^-^ monocytes and KLRG1^+^ NK cells in *Duoxa2*^-/-^ mice were unanticipated (**Fig 5B**). Thus, 3i offers a foundation for exploring how immune variation may be influenced by *Duoxa2-*dependent endocrine effects.

**Figure 5:**
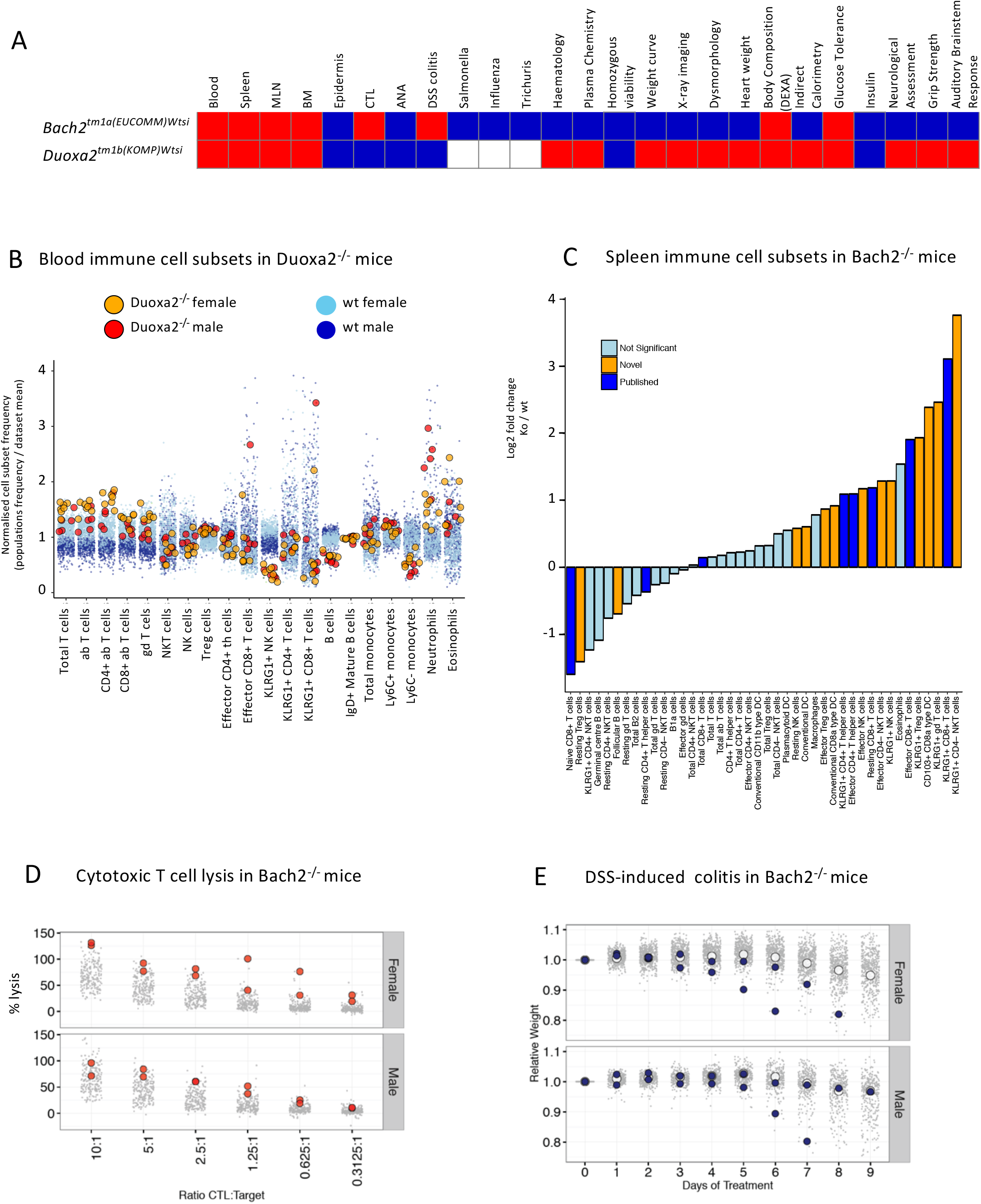
Examples of genes that impact upon the immune system and physiology. A. Overview of two genes that exhibit broad immunophenotypes. Columns on the heat map represent different tests: red, significantly different from wt; blue, not significantly different from wt; white, not performed. B. *Duoxa2^-/-^* mice display multiple phenotypic abnormalities. Relative PBL cell subset frequencies for *Duoxa2^-/-^* (n=11) *versus* wt mice (n>450 per sex). Dark and light blue denote wt males and females; red and orange denote *Duoxa2^-/-^* males and females. C. *Bach2^-/-^* mice are characterized by a marked increase in effector immune cell subsets. Fold-change in SPL cell subset composition in mutant mice (n=6) *versus* wt controls (n=76). Yellow represents significant cellular phenotypes not previously reported, as detected by reference range; dark blue represents significant cellular phenotypes previously reported; light blue represents non-significant differences. D. Cytotoxic T lymphocytes from *Bach2^-/-^* mice (n=4) exhibit increased cytotoxicity compared to wt controls (n>200 per sex): grey points represent individual wt mice; red circles represent individual *Bach2^-/-^* mice. E. *Bach2^-/-^* mice (n=4) show exacerbated weight loss in response to DSS challenge compared to wt controls (n>300 per sex): grey points represent individual wt mice; white circles represent wt mean values; blue circles represent individual *Bach2^-/-^* mice.

Fourthly, *Bach2^-/-^* mice illustrated how 3i could offer broader perspectives for viewing a well-established, disease-related gene. Thus, *BACH2* deficiencies have been widely associated with autoimmunity, inflammation, and allergy, with such associations being commonly attributed to increased effector CD4^+^ and CD8^+^ T cells, as reported for *Bach2^-/-^* mice^39–43^. In addition, however, the 3i platform identified a greatly expanded range of innate and adaptive immune phenotypes of *Bach2^-/-^* mice (orange bars in **Fig 5C**), of which the most significant was a >10-fold increase in KLRG1^+^CD4^-^ NK cells (p=0.000000011; reference range combined with Fisher’s exact test). These findings collectively suggest that some components of *Bach2*-dependent disease might also reflect precocious innate immune cell activation. Further broadening the perspective on *Bach2* were an abnormally high cytolytic activity of *Bach2^-^* CD8^+^ T cells, and a high susceptibility of most *Bach2^-/-^* mice to DSS-colitis (**Fig 5D, E**).

### Genetic regulators of challenge responses

Of 86 phenodeviants affecting steady-state PB, SPL, MLN, BM or epidermal phenotypes, ∼25% also affected responses to challenges, as judged against wt controls assayed contemporaneously. Again, the hits included several sparsely-studied genes (**Fig 6A**, green circles in upper-right quadrant). The fraction of phenodeviants affecting steady-state responses that also affected challenge responses might have been higher if additional challenges and/or assays of response had been employed. However, given the breadth of challenges utilized, and the biologically significant post-challenge outcome measurements (e.g. survival; weight-loss), it is equally likely that genetic redundancy in immune function exceeds that in the baseline immunophenotype. This would predict that genes affecting challenge outcomes should be enriched in those perturbing more than one steady-state parameter, which was indeed the case (**Fig 6A**; larger circles are relatively enriched in upper-right quadrant *versus* lower-right). The prospect of high functional redundancy would ensure that immunocompetence was sustained across a wide range of inter-individual variation in immune cell subsets, as appears to be the case in humans^1–3^.

**Figure 6:**
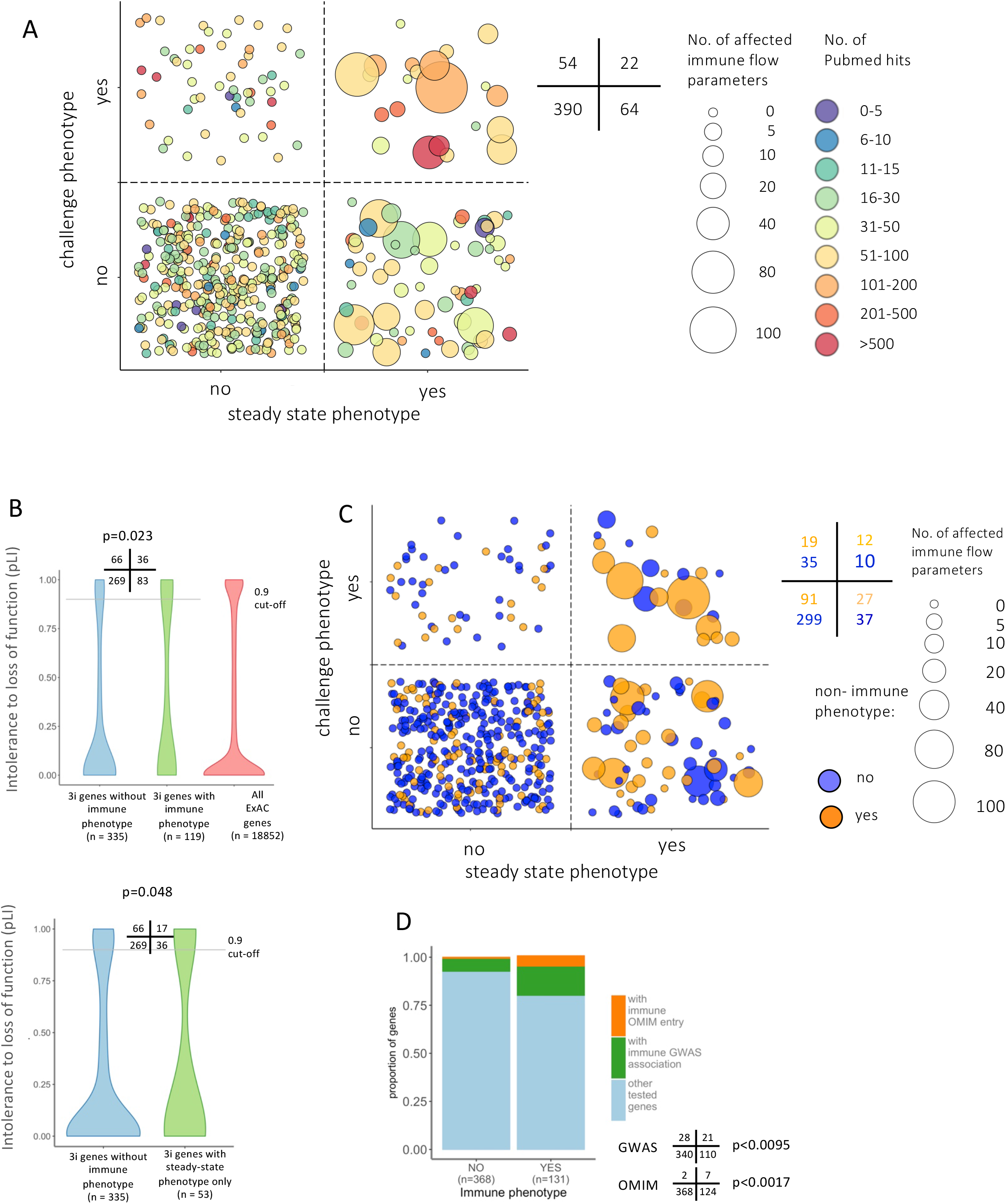
Genes affecting baseline immunophenotypes and challenge responses. A. Each bubble represents one gene; size represents the number of immune flow phenotypes detected for that gene; colour represents the extent to which each gene has been reported on, determined by number of Pubmed citations, as of October 2018. Numbers denote the numbers of genes in quadrants of the figure. B. Human orthologues to murine genes with 3i immunophenotypes are less tolerant to LOF mutations than homologues of 3i genes without immunophenotypes. LOF tolerance scores derived from GnomAD for human orthologues of all 3i genes without hits (blue), for 3i genes with hits (green), and for all GnomAD genes (red). A score of 0 denotes complete tolerance; a score of 1 complete intolerance. Upper panel considers immune hits in all tests (green); the lower graph considers immune hits in steady-state only. In each case, the numbers in quadrants denote the number of genes in the blue and green categories that locate below and above the cut-off, respectively (p-value from Fisher exact test). C. Relationship between immune phenotypes and non-immune phenotypes. Each bubble represents one gene; size represents the number of immune flow phenotypes detected for that gene; colour indicates non-immune phenotype. D. 3i genes with immune hits are more likely to have a known association with disease. Genes with immune-related disease association as defined by OMIM (Online Mendelian Inheritance in Man [OMIM]), or immune-related genome-wide association study (GWAS) are enriched in genes that are 3i hits. Statistical significance was assessed by Fisher’s exact test. Numbers in quadrants represent number of genes with and without OMIM/GWAS association with and without 3i phenotype.

Some phenodeviants, including many sparsely-studied genes, affected functional responses without affecting steady-state immune parameters (**Fig 6A**, upper-left quadrant; see also **Fig 3**). Possibly such gene(s) affect baseline immune parameter(s) not assayed by 3i, while others might reflect gene products which are primarily expressed by non-immune cells and that regulate challenge responses *via* non-immunological mechanisms, e.g. barrier protection. By identifying such genes, the 3i platform offers routes for investigating diverse contributions to host defense.

Given that 57% of the 3i “hits” had never hitherto been implicated in immunobiology, and that the others included many relatively under-studied genes, it was striking that the hits were collectively enriched in genes for which humans show poor tolerance to “loss-of-function” (LOF). Thus, for 454 genes for which human orthologs had been assessed by the Genome Aggregation Database (gnomAD), there was significantly less LOF tolerance for 3i hits (*n*=119) compared either to those genes that did not score as hits (*n*=335) [p=0.023, Fisher’s exact with 0.9 cut-off for LOF^44^], or to all genes based on exomes and genomes from 141,000 individuals^44, 45^ (**Fig 6B**, upper panel). Of note, LOF tolerance was also low for genes that only affected baseline immune parameters (*n*=53) (**Fig 6B**, lower panel), emphasizing the power of a steady-state endophenotype screen such as 3i to identify genes that are functionally relevant to host fitness.

Consistent with this perspective, the 140 immunological phenodeviants were also strikingly enriched in genes that also had non-immunological phenotypes: 58/140 [>41%] *versus* 91/390 [23%] (p=0.000072). Again, this was true for genes affecting baseline immune variation *versus* those that did not: 39/86 [>45%] *versus* 110/444 [25%] (p=0.00021) (**Fig 6C**). Likewise, relative to the genes that did not score, the 131 phenodeviants with identifiable human orthologs were significantly enriched in genes screened because of GWAS implications (21/131 [16%] *versus* 28/368 [7.6%])(**Fig 6D**). This further emphasizes the capacity of steady-state endophenotypes to inform disease associations, evoking the profound utility of increased B cell counts and reduced monocyte numbers in linking a specific *BAFF* gene variant to multiple sclerosis and lupus^15^. Discrete endophenotypes were likewise scored for seven of nine genes with pre-assigned OMIM immunology-related designation (**Fig 6D**).

In sum, through the identification of baseline and/or post-challenge monogenic endophenotypes, the 3i screen has greatly expanded the number and scope of genes implicated in immunobiology. Moreover, given that ∼70% of genes screened were little-studied, it was striking that the 3i endophenotypic screen produced an unbiased enrichment for genes likely to be of human disease-relevance given their low LOF tolerance and/or impacts on general physiologic traits.

### Genetic regulation of correlation networks

The 3i data-sets were next used to ask whether genes affecting one or more steady-state immune parameter(s) might segregate according to whether or not they also disrupted the correlations connecting immune parameters: i.e. whether or not they impacted upon immunological structure. **Fig 7A** (left panel) schematises hypothetical distribution clouds of data-points for three phenodeviants (yellow, pink, green) displaying extreme values for cell population B. Of note, whereas the yellow and green genes maintained proportionality with population A, the pink gene broke that correlation.

**Figure 7:**
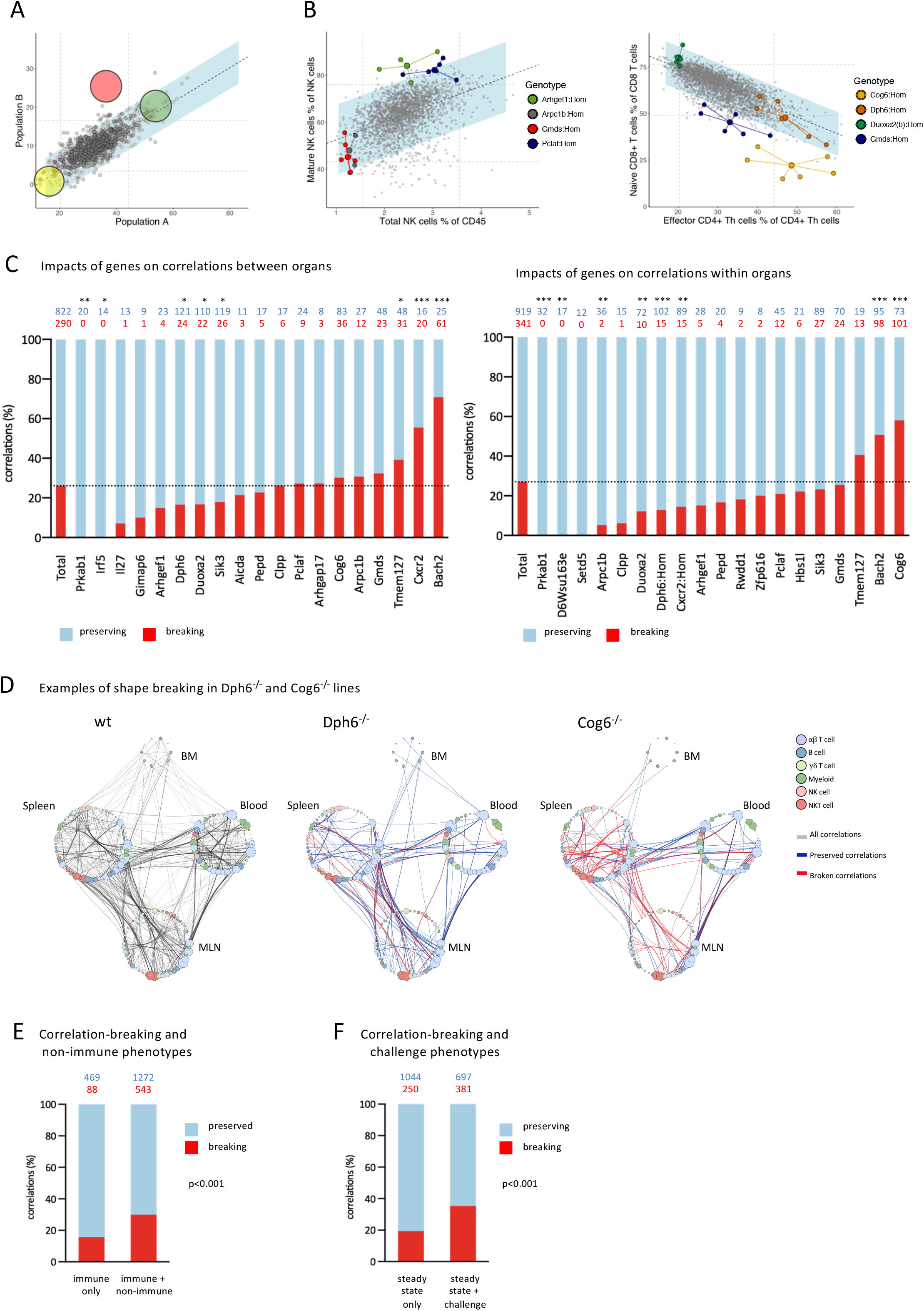
Phenodeviants preserve or break immunological structures. A. A schematic illustrating the concept of structural perturbation. The yellow and green genes are theoretical hits in both correlated parameters, and exist as an exaggeration of the normal relationship that exists at steady state (grey dots represent wt mice). The pink gene is also a hit in both parameters, but breaks the correlation by falling outside the blue corridor that represents a 95% prediction confidence interval around the correlation line. B. Examples of genes that preserve or disrupt immunological structure. Plotted are data frequencies of SPL subsets (indicated on x- and y-axes) as determined by flow cytometry. Data from mutant lines with phenotypes are highlighted in different colours; large dots represent the x/y centroid (mean) values, small dots represent data points for each mouse. Grey data points represent wt mice and mutant mice that are comparable to wt mice for both parameters shown. C. Genes differ in their capacity to break or preserve relationships. Stated in red font are the number of correlations that each cited gene disrupts and in blue font the residual number of correlations that each cited gene preserves, the total being all correlations contributed to by the steady-state parameters that the cited gene affects. These are represented as percentages in the left-hand graph (correlations across organs) and right-hand graph (correlations within the same organ). Only genes affecting parameters which contribute to >10 correlations are depicted. Statistical significance was determined by Fisher’s exact test in comparison to the data set average (dotted line). *p<0.05, **p<0.01, ***p<0.001 D. Examples of genes that either preserve or disrupt many correlations. Plotted are parameters (colour-coded according to cell lineage) in BM, SPL, MLN, and PBL. and the correlations that link them in wt mice (left panel, grey lines) or in *Dph6^-/-^* mice (middle) or *Cog6^-/-^* mice (right), in which cases blue and red lines denote correlations that are preserved or broken, respectively. E. Genes that score in non-immune tests and immune tests collectively break more steady-state immune correlations than do genes that have hits only in immune parameters (significance established by Probit regression). F. Genes that score in challenge assays and steady-state immune tests collectively break more steady-state immune correlations than do those that score only in steady-state immune parameters (significance established by Probit regression).

Real examples of this are shown in **Fig 7B**: *Gmds* and *Arpc1b* mutants displayed very few NK cells but maintained proportionality with mature NK cells, whereas most *Arhgef1^-/-^* and *Pclaf1^-/-^* mice sat outside the confidence limits of the relationship between total and mature NK cells, showing disproportionately high NK maturation. Likewise, *Duxoa2^-/-^* and *Dph6^-/-^* mice, whose effector CD4^+^ T cell numbers were atypically low and high, respectively, each preserved an inverse relationship of effector CD4^+^ Th cells and naïve CD8^+^ T cells, whereas this correlation was broken by deficiencies in *Gmds* and *Cog6*. Hence, correlations that confer structure on the immune system can be resilient to one genetic perturbation, but be broken by a different one.

Overall, we observed that genes with the potential to affect ≥10 correlates collectively preserved ∼75% of intra- and inter-organ correlations (**Fig 7C**; “Total”). Hence, the overriding impression was that immune correlations were relatively resilient to genetic perturbation of discrete immune parameters that can underpin immune variation. Nonetheless, it was inevitably the case that mutations of some genes that affected large numbers of immune parameters were more likely to disrupt substantially more correlations, as was evident for *Cog6* and *Bach2* (**Fig 7C**). This notwithstanding, structural breakage did not simply reflect the number of immunophenotypes affected by any one gene. Thus, most intra-organ and inter-organ correlations of immune cell subsets were preserved in *Dph6^-/-^* mice which displayed a similar number of steady-state immune parameters (e.g. altered cell subsets) to *Cog6^-/-^* mice: for example, compare the density of red lines (broken correlations) *versus* blue lines (preserved correlations) in the spleens of the two strains in **Fig7D**. This argues that specific genes underscore the existence of certain correlations and hence underpin immunological structure. Added to this, *Arpc1b* broke many inter-organ correlations while preserving most intra-organ correlations (**Fig 7C**), indicating that different genes regulate immune subset relationships in different ways.

Given that many immune parameters also correlated with specific physiologic measures, thereby composing structures of the kind illustrated in Fig 2C, it was appropriate to ask whether genes with non-immunological phenotypes were more likely to disrupt immunological structure. Indeed, baseline structure seemed much less resilient when discrete physiological traits were also perturbed, with such genes collectively breaking significantly more correlations than were broken by genes only affecting immune parameters: 30% *versus* 16% (p<0.001, Probit regression) (**Fig 7E**). Additionally, it was noteworthy that genes which impacted upon challenge responses as well as upon the baseline immunophenotype broke significantly more correlations than did genes affecting only the baseline immunophenotype: 35% *versus* 19% (p<0.001, Probit regression) (**Fig 7F**).

The results shown in Figs7E and 7F were robust to controlling for unequal numbers of correlations per gene (non-immune p<0.001, challenge p<0.001) and to allowing for any dependencies of correlations within a gene by using cluster-robust standard errors (non-immune p=0.007; challenge p=0.021). Moreover, controlling for the influence of non-immune traits and challenge responses in the same regression analysis confirmed separate contributions of non-immune and challenge phenotypes (p<0.001 for non-immune controlling for challenge; p<0.001 for challenge controlling for non-immune). Clearly, the association of defects in immunocompetence with compromises to structural integrity in the baseline immune system strongly suggests the importance of immunological structure and makes a compelling case for measuring immune subset correlations as a means to monitor functionally significant immune variation.

## Discussion

The development and implementation of the 3i platform has combined genetic screening and immunophenotyping at scale. Thereby, it has established that variation in the immunophenotype can be significantly shaped by genetics, substantially expanding the number and diversity of genes shown to have monogenic impacts on steady-state immune parameters and/or challenge responses. Those findings compose a resource for generating hypotheses to underpin subsequent in-depth studies. The potential value of this is highlighted by recent evidence that baseline endophenotypes can critically aid the causal association of genetic variants with human disease^15^. In this vein, it is noteworthy that 3i’s screening mostly of little-studied genes conspicuously enriched for those for which humans show poor tolerance to loss-of-function. Their individual follow-up may expose understudied immunoregulatory pathways that can inform polygenic human diseases where single-gene effects sizes are small. Those pathways will probably include those that act immune cell autonomously and those that act *in trans*.

At the same time, the scale and robustness of 3i, including its minimization of technical and batch variation, has provided insight into the nature and scope of immune variation in one of the community’s most-studied animal model, the C57.BL/6N mouse, with comprehensive, in-depth data-sets curated on a sustainable, user-friendly web interface www.immunophenotype.org. In practical terms, the 3i platform also offers protocols by which to quickly obtain broad immunophenotypic data from mice in myriad experimental scenarios. In this regard, it is noteworthy that the relative proportions of cells composing the spleen were recently shown to be as effective an indicator of autoimmune dysregulation as the cells’ micro-anatomical localization^46^.

The scale and throughput of the 3i screen yielded substantial data-sets supporting the view that the baseline immune system is strongly sexually dimorphic^47, 48^, but showed that this applied highly specifically. These findings should be accommodated in the design of mouse experiments, and whilst the specifics of sexual dimorphism will differ from mouse to human^48^, the findings should likewise be considered in the design of immune interventions and immune-monitoring strategies in humans.

The scale, throughput, and robustness of the 3i screen were also sufficient to reveal dense networks of correlations of immune parameters with themselves and with general physiologic traits. Such correlations will *de facto* impose structures constraining immune variation. Although high-level structure was more evident in males, we believe that similar structures exist in females but is masked by the higher variability in general physiology. Consistent with this, males and females showed comparable frequencies of correlations between immunological parameters.

The correlations charactering the baseline immunophenotypes of male and female C57.BL/6N differ in detail across different organs, and they will inevitably differ across mouse strains and across species. This is not to diminish the importance of the correlations revealed by 3i, since they provide a structural framework for viewing immune variation more generally. For example, immunological structures may sustain balanced immune interactions that to some degree insulate immune function from inter-individual and sexually dimorphic variation. Likewise, the co-ordination of immune cell subsets reflected by immunologic structure may confer agility on the immune system to respond to diverse challenges. This would be entirely consistent with the observation that most phenodeviants in the steady-state failed to impact the challenge responses as measured by 3i, whereas correlations between immune parameters (i.e. the underpinnings of immunological structure) were more frequently disrupted by genes that impaired challenge response(s).

This association of immunological structures with biologically significant outcomes makes a compelling case for including structural metrics (e.g. subset ratios *versus* consensus values) in immune-monitoring strategies. In support of this, *Bach2*, *Cog6* and *Arpc1b* which are strongly associated with human disease, were readily scored by the criterion of disrupting of inter-organ correlations^39–41, 49–51^. Moreover, therapeutic responsiveness to checkpoint blockade was recently associated with a ratio of CD8^+^ T cell subtypes, rather than with subsets viewed individually ^52^.

Likewise, immune-monitoring strategies might usefully include general physiologic traits, since correlations linking immune parameters were more frequently disrupted by genes that also affected non-immunological parameters. Moreover, while many will vary, some of the core correlations may be conserved cross-species. For example, the numerous correlations with HDL-cholesterol may be relevant to the established anti-inflammatory impacts of HDL, that have to date been primarily attributed to the down-regulation of TLR-mediated cytokine induction^53^. Likewise, the many immunological correlations with Rdw were unanticipated but are potentially linked to the recently emerging strength of Rdw as a predictor of all-cause mortality, perhaps reflecting its association with pathologic inflammation germane to cardiovascular disease and cancer^30^. Collectively, the robust correlations revealed by 3i emphasize the need for more intensive studies of how immunology is integrated with specific components of general physiology.

Finally, the density of correlation networks revealed by 3i makes the case for investigating their biological basis. Do they, for example, reflect gene regulatory networks common to the homeostasis of different cell populations, and/or robust intercellular circuitry, as was recently illustrated for macrophages and fibroblasts?^23^ Such processes may themselves be regulated by factors such as diet and hormones, reflected by the correlations of immune parameters with discrete physiologic traits. The biological basis of correlation networks may create foundations for better understanding how mutations in specific genes, e.g. *Bach2*, and/or in the super-enhancers that control those genes may provoke severe immunopathology because of their impact on immunological structure^43^.

## Author contributions

Conceptualisation: J.W., F.P., G.D., W.J., C.L., R.J.C., K.M., G.G., R.G., D.A., A.C.H. Methodology: L.A-D., A.G.L., A.L., D.U., S.C., A.O.S., N.Sa., M.D., K.R.B., B.M., J.I. P.B., K.I.H., E.C., S.F., T.L.C., B.W., A.R., S.D., J.M., A.Y., M.L., G.S-Z., A.C., R.B., G.D., W.J., C.L., R.J.C., K.M., G.G., R.G., D.A., A.C.H. Software: A.L., M.G., N.R., N.A.K., D.M., A.R., S.D., J.M., A.Y., M.L., J.A., R.B. Website: L.A-D., A.L., K.O.B., J.W., J.M., A.T.M., T.M. Validation: L.A-D., A.G.L., A.L., D.U., S.C., A.O.S., N.Sa., M.D., K.R.B., B.M., J.I., P.B., E.C. Formal analysis: L.A-D., A.G.L., A.L., D.U., S.C., A.O.S., N.Sa., M.D., K.R.B., B.M., J.I., P.B., E.C., A.C.H. Investigation: L.A-D., A.G.L., D.U., S.C., A.O.S., N.Sa., M.D., K.R.B., B.M., J.I. P.B., K.I.H., E.C., S.F., T.L.C., H.W., L.K., K.H., C.B., G.N., E.C., B.W., G.S-Z., A.C., C.B.R., A.P-K., M.E., N.St., M.P. Data curation: L.A-D, A.L. Writing – original draft: A.C.H. Writing – Review and editing: L.A-D., A.G.L., A.L., D.U., A.O.S., N.Sa, N.A.K., J.A., G.D., R.J.C., G.G., R.G., D.A., A.C.H. Vizualisation: L.A-D., A.G.L., A.L., A.C.H. Supervision: R.B., G.D., W.J., C.L., R.J.C., K.M., G.G., R.G., D.A., A.C.H. Project Administration: L.A-D., J.W., S.C., A.O.S., R.R-S., A.C.H. Funding Acquisition: R.B., F.P., G.D., W.J., C.L., R.J.C., K.M., G.G., R.G., D.A., A.C.H.

## Acknowledgements

We thank many colleagues for advice, particularly Drs S. Heck of the Biomedical Research Centre of Guy’s and St Thomas’ Hospital and King’s College London; D. Davies of the Francis Crick Institute; F. Geissmann, J. Strid and O. Sobolev in the early stages of programme planning; and N. Karamanis and K. Hawkins for UX testing of the website. Supported by grants 100156/Z/12/Z and from the Wellcome Trust, and by grants and facilities provided to A.C.H. by the Francis Crick Institute, which receives its core funding from Cancer Research UK (FC001093), the UK Medical Research Council (FC001093), and the Wellcome Trust (FC001093). KB, TM, JW were supported by the NIH Common Fund [UM1 HG006370] The development of automated flow analysis in R.B.’s group was supported by Genome BC (SOF152), NSERC, International Society for Advancement of Cytometry Genome Canada, Genome BC (252FLO) and CIHR.

## Materials and Methods

### Contact for Reagent and Resource Sharing

For additional information about reagents and resources, contact Adrian Hayday at.

### Experimental design

All assays relied on the fastidious application to cells and tissues of intensively piloted, robust, optimised Standard Operating Procedures (SOPs) employing high-resolution, quantitative protocols, whose high reproducibility was monitored over time (e.g. **Fig S1F**).

Mice were randomly allocated to the experimental groups (wt *versus* ko) by Mendelian inheritance. The experimental unit in the study was the individual mouse. For the majority of tests, operators were blinded with regard to the genetic identities of mice. Further detailed experimental design information is captured by a standardized ontology as described^19^, and is available from the IMPC portal <http://www.mousephenotype.org>. The steady-state screens integrated within the HTS followed a multi-batch design, in which a baseline set of control data was constantly collected by phenotyping wt mice of both sexes along with mutant mice at least once per week. As soon as mutant mice from the breeding colonies reached appropriate ages, they were issued to the pipeline until sufficient number of males and females of each genotype were assessed. In this way, animals from each mutant strain were tested on two to five days, interspersed throughout experiment duration, rather than within one batch. For advantages and robustness of such design in an HTS see ^21, 56–58^. The challenge screens implemented a parallel group study design.

Compliant with HTS practice, the numbers of mice examined in each assay were dictated by costs, logistics, and power, with bespoke statistical tests (see below) applied to identify genes (so-called “hits”) perturbing defined components of the immune system. Two to seven homozygote mice of each sex per mutant line were assessed per test. If no homozygotes were obtained from ≥28 offspring of heterozygote intercrosses, the line was deemed homozygous lethal. Similarly, if <13% of the pups resulting from intercrossing were homozygous, the line was judged as being homozygous subviable. In either case, heterozygotes were phenotyped. The numbers and sex of animals tested per genotype and assay are summarized in **Table S3**.

### Ethical compliance

Mouse use in this study was justified based on their facilitating a large variety of phenotypic tests to be carried out on a sufficient number of individuals in a controlled environment. The care and use of mice in the study was conducted in accordance with UK Home Office regulations, UK Animals (Scientific Procedures) Act of 2013 under two UK Home Office licenses which approved this work (80/2076 and 80/2485) which were reviewed regularly by the WTSI Animal Welfare and Ethical Review Board.

All efforts were made to minimize suffering by considerate housing and husbandry, the details of which are available at the IMPC portal: http://www.mousephenotype.org/about-impc/arrive-guidelines. Animal welfare was assessed routinely for all mice involved. Adult mice were killed by terminal anaesthesia followed by exsanguination, and death was confirmed by either cervical dislocation or cessation of circulation.

### Animals

Mice were maintained in a specific pathogen-free unit under a 12-hour light, 12-hour dark cycle with water and food *ad libitum* (Mouse Breeders Diet (LabDiets 5021-3, IPS, Richmond, USA), unless stated otherwise. Mice where housed in Tecniplast Sealsafe 1284L (overall dimensions of caging (L × W × H): 365 × 207 × 140 mm; floor area = 530 cm^2^) at a density of 3-5 animals per cage, and provided with a sterilized aspen bedding substrate and cardboard tubes and nestlets for environmental enrichment.

### Mutant mouse production

Mice carrying knockout first conditional-ready alleles were generated on a C57BL/6N background using the EUCOMM/KOMP Embryonic Stem (ES) cell resource, with ES quality control and molecular characterization of mutant mouse strains performed as described previously ^59^. Upon completion of phenotyping, genotyping was repeated and data were only accepted from mice for which the second genotype was concordant with the original. The knock-in first strategy first generates mice that still possess the full sequence of the targeted gene, interrupted in a crucial exon by the inserted cassette. These tm1a alleles result in functional knockout lines in most cases, but can carry residual expression of the targeted gene. These tm1a alleles can be converted into unequivocal full knockout tm1b alleles by excising the inserted cassette with the targeted essential exon. Details of alleles used can be found in Skarnes *et al*^19^. All lines are available from www.knockoutmouse.org or.

### Non-immune phenotyping

Non-immune phenotyping (summarized in **Fig S1A**) was conducted as described ^21^. Screening for phenotypes in *Citrobacter* infection was replaced by the 3i challenges.

### Immunological steady-state phenotyping

Tests conducted in the steady-state (PBL, SPL, MLN, BM, ear epidermis, antinuclear antibodies, cytotoxic T lymphocytes function) were conducted on the same 16-week-old mice that were subject to broad non-immunological phenotyping procedures^21^. Non-fasted mice were terminally anaesthetised using Ketamine (100 mg/kg)/Xylazine (10 mg/kg) injection. Organs were harvested and either analysed directly (PBL) or shipped in HBSS at 4°C for analysis on the same day off-site. Readouts of the respective tests and numbers of mice used are summarized in **Fig S1** and **Table S3**.

#### Single cell preparation of immune cells from spleen

After removing the fat, spleens were transferred into Miltenyi C-tubes with 3 ml of enzyme buffer (PBS Ca^2+/^Mg^2+^, 2% FCS (v/v), 10 mM HEPES, Collagenase (1 mg/ml), and DNAse (0.1 mg/ml). Samples were then processed using a Miltenyi Gentle MACS dissociator (SPL program 01) and incubated at 37°C for 30 minutes. After the incubation, samples were processed again in the Miltenyi Gentle MACS dissociator (SPL program 02) and the enzyme reaction was stopped adding 300 μl of stop buffer (PBS, 0.1 M EDTA). Samples were filtered through 30 μm filters and centrifuged for 5 minutes at 400 x g at 8°C. The pellet was resuspended in FACS buffer (PBS, Ca^2-/^Mg^2-^, 0.5% FCS, EDTA 2mM), incubated for 60 seconds in RBC lysis buffer and then washed with FACS buffer. Samples were transferred to 96-well V bottom plates and incubated in 50 μl of RBC lysis buffer (eBioscience) for 1 minute prior to antibody staining.

#### Single cell preparation of immune cells from mesenteric lymph nodes (MLN)

After removing the fat, MLN were transferred into 1.7 ml microfuge tubes with 200 μl of buffer (PBS Ca^2+/^Mg^2+^, 2% FCS (v/v), 10 mM HEPES) and ruptured using small plastic pestles. 400 μl enzyme buffer (PBS Ca^2+/^Mg^2+^, 2% FCS (v/v), 10 mM HEPES), Collagenase (1 mg/ml), and DNAse (0.1 mg/ml) were and samples were incubated for 15 minutes at 37°C. The reaction was stopped by adding 60 μl stop buffer (PBS 0.1 M EDTA) and samples filtered through 50 μm filters and centrifuged for 5 minutes at 400 x g at 8°C. The cell pellet was resuspended in FACS buffer (PBS Ca^2-/^Mg^2-^, 0.5% FCS, EDTA 2 mM) and transferred to 96-well V bottom plates for antibody staining.

#### Single cell preparation of immune cells from bone marrow (BM)

The tibia was cleared of muscle tissue and cut below the knee and above the ankle. The open bone was placed into a cut pipette tip placed into a microfuge tube, thereby keeping the bone away from the bottom of the tube and allowing the bone marrow to be centrifuged out of the bone at 1000 *×* g for 1 minute. The bone was discarded and the pellet resuspended in 50 μl RBC lysis buffer at room temperature for 1 minute. Cells were washed in 200 μl FACS buffer and again centrifuged at 1000 *×* g for 1 minute. Cells were resuspended in 400 μl FACS buffer and transferred to 96-well V bottom plates for antibody staining.

#### Blood preparation

Blood was collected into EDTA coated tubes (peripheral blood leukocytes assay) *via* the retro-orbital sinus. Whole blood for peripheral blood leukocyte assays was stained with two titrated cocktails of antibodies (**Table S1**). Using the white blood cell count obtained from the haematological analysis, absolute cell counts were derived for each population and reported as cells/µl.

### Immunophenotyping by flow cytometry

Single cell suspensions were incubated with Fc-block for 10 minutes at room temperature, washed four times, first with FACS buffer and then with PBS, and then incubated with live/dead ZiR dye (BioLegend) for 10 minutes at room temperature. Samples were washed again with FACS buffer and incubated with antibody cocktails (see **Table S1**) at 4°C for 20 minutes. Samples were washed twice with FACS buffer and measured on a BD Fortessa X-20 equipped with 405 nm, 488 nm 561 nm and 644 nm lasers (see **Table S1**). Full details of instrument setup are available at www.immunophenotyping.org. Panels were modified slightly in summer 2014 in order to better correspond to the IMPC panels (T cell panel: 9^th^ June 2014, B cell panel: 15^th^ September 2014). Data before and after this data split were analysed separately. Data were analysed with FACSDiva and Flowjo software. FCS files are available from flowrepository.org.

#### Flow cytometry quantification (SPL, MLN and BM)

Additionally to the manual gating performed at the time of data acquisition, collected flow cytometry data for SPL, MLN and BM were gated computationally using flowClean, UFO, and flowDensity^22, 60^. FlowClean was used to perform acquisition-based quality checking to remove anomalous events. Files with fewer than 20,000 events were then removed from further analysis. UFO was used to identify outlier samples (e.g., batch effects). FlowDensity was used to enumerate cell populations by automating a predefined gating approach using sequential, supervised bivariate gating analysis to set the best cut-off for an individual marker using characteristics of the density distribution. The parameters for each individual cell population were pre-defined once for all files. The automated analysis data was validated against matched manually analysed data. Gating strategies are outlined in **Fig S1B** and in **Fig S4 A-D**). Assessment of absolute cell counts were not compatible with the high-throughput workflow employed in the study for SPL, MLN and BM. All cell subset frequencies are presented as a percentages of a relevant parent populations.

### Ear epidermis immunophenotyping

Epidermal sheets from mouse ears were treated with hair removal cream (Nair) for 4 minutes at room temperature. After removing the cream by extensive washing in PBS, ears were split into dorsal and ventral sides and incubated dermal side down for 35 minutes at 37°C in 0.5 M ammonium thiocyanate (Sigma-Aldrich). Epidermal sheets were gently peeled from the dermis in PBS, and fixed in cold acetone for 20 minutes at −20°C. After washing in PBS, epidermal sheets were incubated for 1 hour at room temperature with 5% (wt/vol) FCS in PBS and were stained for 1 hour at 37°C with Vγ5 TCR-FITC (clone 536; BD), MHCII-AF647 (I-A/IE; BioLegend) and CD45-eFluor450 (eBioscience). Epidermal sheets were washed extensively in PBS and mounted on slides with ProLong Gold antifade medium (Life Signalling Technologies). Leica SP2 or SP5 confocal microscopes equipped with 40 x 1.25 NA oil immersion lens and 405 nm, 488 nm and 633 nm lasers were used to record 1024 x 1024 pixel confocal z-stacks with Leica Acquisition Software. The confocal records were processed and quantified with Definiens Developer XD^R^ software using a custom made automated protocol where images were smoothed with a sliding window Gaussian pixel filter, segmented by an automated Otsu’s method and then filtered based on object size and morphology parameters to detect cells in each fluorescence channel. Further, in order to quantify the number and length of dendrites, the detected cells were skeletonised in 3D to determine the points where dendrites start to branch out of the cell body.

### Antinuclear Antibody Immunophenotyping

Murine serum samples were obtained and stored at −20**°**C prior to analysis (after dilution 1:100 in PBS) by incubating on commercially-sourced substrate slides (A. Menarini Diagnostics Ltd.) coated with HEp-2 cells for 30 minutes at room temperature in a humidifying tray. Samples were removed and slides were washed twice with PBS for 5 minutes and once with water for 5 seconds. Slides were incubated with FITC-conjugated goat anti-mouse IgG, diluted 1:500 and incubated for 20 minutes at room temperature in a humidifying tray in the dark. The secondary antibody was removed and slides washed twice with PBS for 5 minutes and once with water for 5 seconds, both in the dark. Slides were mounted in medium and stored at 4**°**C prior to imaging for 400 ms at 20 x magnification in the GFP channel on a Nikon wide-field TE2000U Microscope or a Deltavision Elite widefield system based on an Olympus microscope. Images were subject to multi-parametric analysis in Fiji. Samples were scored from 0-4 according to intensity based on control samples and commercially sourced FITC QC beads. Samples scored ≥2 were marked as ANA positive. All sera flagged by automated image analysis as putative positives *vis-à-vis* a contemporaneous standard were manually cross-checked for *bona fide* nuclear localization before scoring.

### Cytotoxic T Lymphocyte Immunophenotyping

Mouse splenocytes were isolated using 70 μM cell strainers (BD Plastipak) and cultured in T cell media (TCM: 500 ml RPMI, 500 μl B-ME, 5 ml NaPyr, 5 ml pen/strep, 5 ml L-glut, 50 ml 10% heat inactivated FCS and 100 μl IL-2) on 6-well plates pre-coated with 0.5 μg/ml anti-CD3*ε* antibody and 1 μg/ml anti-CD28 antibody (1.7*×*10^6^ cells/well) for 48 hours. Plates were washed and cells where cultured for a further 8 days with daily passage prior CTL assay.

Cytotoxicity assays were performed using a CytoTox96 Non-Radioactive Cytotoxicity Assay kit (Promega UK Ltd). Cells were washed and re-suspended in killing assay media (KAM: 500 ml RPMI-phenol red, 10 ml 10% heat inactivated FCS and 5ml pen/strep), with CTLs at a concentration of 0.1 × 10^6^ and P815 target cells at a concentration of 0.1 × 10^6^. Purified hamster anti-mouse CD3*ε* antibody was added to P815 target cells.

P815 cells were added to a serial dilution of CTL samples and incubated for 3 hours. Lysis buffer (Promega UK Ltd.) was added to control samples and incubated for 45 minutes. Supernatants were harvested and substrate mix (Promega UK Ltd) was added prior to a 30-minute incubation in the dark. Stop solution (Promega Corporation UK Ltd) was added to halt the reaction and results acquired using a spectrophotometer (VersaMax, molecular devices).

Flow cytometric analysis was performed to assess the percentage of CD4^+^ and CD8^+^ cells within the cell culture. The cell suspension was washed in FACS buffer and the cell pellet re-suspended in a staining master mix (FACS buffer solution + 1:200 anti-CD8*α* APC and 1:200 anti-CD4 PE). Tubes were then incubated in the dark for 7 minutes at room temperature before the antibody was washed off and cells resuspended in FACS buffer. Results were acquired on a FACS Calibur machine and analysed using FlowJo 10 software.

### Challenge screens

Challenge screens were conducted on separate cohorts of mice from the same breeding colony used for the steady-state screens.

#### DSS colitis challenge

Colitis was induced by adding 1.5% (w/v) DSS (Affymetrix, Inc.) to drinking water for 7 days, followed by 3 days with regular drinking water, in animals aged between 5 and 18 weeks (mean age 9 weeks). Mice were weighed every day and culled if weight loss reached 20% of starting weight.

For histological assessment of intestinal inflammation, mice were sacrificed at day 10 by cervical dislocation, and samples from mid and distal colon taken. Tissue sections were fixed in buffered 10% formalin; paraffin-embedded; cut; and stained with haematoxylin and eosin. Colon histopathology was blind-graded semi-quantitatively on a scale from zero to three, for four criteria: (1) degree of epithelial hyperplasia/damage and goblet cell depletion, (2) leukocyte infiltration in lamina propria, (3) area of tissue affected, and (4) presence of markers of severe inflammation, including crypt abscesses, submucosal inflammation, and oedema. Scores for individual criteria were added for an overall inflammation score of between zero and twelve for each sample. Scores from mid and distal colon were then averaged to obtain inflammation scores for each mouse colon.

#### Salmonella typhimurium challenge

Groups of 8 mutant and 8 C57BL/6N wild type mice were challenged intravenously with 5 x 10^5^ colony forming units (cfu) *Salmonella typhimurium* M525 :: TetC, (Fragment C of tetanus toxin, to act as an antigen for subsequent antibody quantification), and followed for 28 days. On day 14 post-infection (pi), four mutant and four wt mice were sacrificed by cervical dislocation and organs (spleen, liver and caecum) removed. A small piece of spleen and liver was fixed in 4% formalin and then later processed to paraffin blocks as an infected tissue biobank. The rest of the organs were weighed then homogenized, serially diluted and plated to determine viable bacterial load. At day 28 pi, the remaining four mice were culled by cardiac puncture under terminal anesthesia and organs removed and processed, as above. The blood was allowed to clot, then centrifuged, serum collected and used to detect TetC antigen specific antibodies by enzyme-linked immunosorbent assay (ELISA). Mice were weighed and monitored daily for signs of pathophysiology.

#### Influenza challenge

Mutant and wt mice (5-21 weeks of age) were lightly anesthetised and intranasally inoculated with 171 or 227 p.f.u. of A/X-31 (H3N2) influenza in 50 μl of sterile PBS. Mouse weight was recorded daily and the percent reduction was calculated from their weight on day 0. Mice were sacrificed by cervical dislocation on day 10 pi, and the area under the curve from day 0 to 9 pi was calculated. Mice exceeding 25% total weight loss were culled in accordance with UK Home Office guidelines.

#### Trichuris muris challenge

The Edinburgh (E) strain of *Trichuris muris* was used in all experiments. Female mice (6-12 weeks old) were orally infected with 400 embryonated eggs. Mice were culled by cervical dislocation at day 32 pi, blood was collected by cardiac puncture for serum recovery, and caecum/proximal colon was dissected to inspect for worm presence by stereomicroscope. Levels of parasite-specific serum IgG1 and IgG2a Ab were by ELISA: briefly, ELISA plates were coated with *T. muris* excretory/secretory (E/S) antigen at 5 μg/ml. Serum was diluted 1/40, and parasite-specific IgG1, IgG2a and IgE detected with biotinylated anti-mouse IgG1 (Biorad), biotinylated anti-mouse IgG2a (BD PharMingen), and anti-mouse IgE (BioLegend), respectively. To generate *T. muris* E/S antigen, live adult worms were incubated at 37°C for 24 hours in RPMI-1640 medium (Gibco, UK) supplemented with 500 U/ml penicillin, 500 μg/ml streptomycin and 2 mM L-glutamine (all Gibco, UK). Supernatants were removed, centrifuged at 2000 x g for 15 minutes, filtered through a 0.22 μm filter (Millipore, UK), concentrated using a 10 kD molecular weight cut off Centriprep concentrator (Amicon, UK) and dialysed against PBS over a 24-hour period. The supernatant was subsequently filtered again and protein concentration determined before use.

### Statistical analysis

Sample sizes for each experimental group are included in the figure legends and **Table S3**. Measures were taken from distinct samples in almost all experiments. Exceptions are body weight, where the weight of the same animal was recorded repeatedly during the course of the experiment and the CTL assay, in which T cells from the same culture were used for different effector : target ratios. Tests for all assays are listed in **Table S3**. Reference-range tests which require no assumptions were used for most assays. Adjustments for multiple comparisons were not made unless stated in test description, as individual comparisons were not independent (e.g. percentages for different cell types determined by flow cytometry are often nested and dependent on each other). Low false positive rates were instead ensured by using conservative reference range cut-offs and monitoring false positive rates by simulation (**Fig S3**). All tests used were two-sided. Co-variates were included in the analysis indirectly, by dividing animals into sex-specific groups or matching ko and wt by assay date/weight/age (MMR analysis described below) and in the analysis presented in figures 7 E and F where covariates tested are described in the text. A description of statistical parameters is included into the figure legends. Effect sizes are given when fold change between sexes was analysed. All significant calls and borderline candidates were manually reviewed by biological experts.

Experimental batch effects were minimised by the experimental design, the statistical analysis and by expert review. Experimental set-up: Mice were phenotyped on at least two different days and typically three to four different days. Statistical analysis: Stringent reference range hit calling required ≥60% of samples to be in the upper 2.5% or lower 2.5% of the w.t. distribution. The influence of temporal drift in flowcytometric data was minimized by using as controls only the 70 wt mice that were closest in time for any given parameter. Reference ranges have been shown by the WTSI team to be stable for ≥60 mice. Expert review: Experts reviewed all hits and borderline candidates and excluded candidates that suffered from batch effects despite the measures taken above.

Coefficient of variation (CV) was estimated as a ratio of the standard deviation of the wt population to its mean (for both sexes together, unless indicated otherwise). Note that since a data-set is expressed as frequency of parent, it is bound and CV estimates are less accurate close to the boundary.

#### Sexual dimorphism

Significance of sex as a source of variation in wt animals was tested in a mixed model (sex as explanatory variable, assay date as a random effect) by examining the contribution of sex to the model and whether the variance was homogeneous between sexes^61^. The effect of multiple testing was managed with Bonferroni correction to control the family-wise error rate to 5%. For Principal Component Analysis and Linear discriminant analysis (LDA), only wt samples with a complete set of flow parameters were used. Data was scaled. Accuracy of classification by sex in LDA was checked with leave-one-out cross-validation.

#### Estimation of false positive rates

A mutant mouse line was mimicked by randomly selecting N wt animals (N depending on the screen) and assessed whether it would be called a hit using the RR approach. To reflect the data structure and address the potentially confounding batch effect of test days, the number of experimental days from which animals were drawn and their sex was set according to the distributions observed across all tested mutant lines. For example, in the lab, bone marrow was collected on five different days in 36% of mouse strains; in 38% on four different dates; in 11% from three different days, etc. When strains were tested on four assay days, in 78% of cases females were tested on two days, etc. The same data structure was imposed on the draws for the false positive rates. Thereupon, the same rubric used for calling hits from real data was applied, except that the expert data review step (see above) was replaced by filtering out samples from the days when the median of wt animals and non-significant mutant strain samples was further than two median absolute deviations (MADs) from the overall median in wt animals. 10,000 draws were performed for each parameter to determine the rate at which false positives would occur.

#### Correlation analysis

Pearson correlations between parameters in the flow cytometry screens as well as between the flow cytometry screens and non-immune screens were identified in wt mice for pairwise complete observations, accumulated throughout whole duration of the experiment. Correlations with parent and sister populations were excluded from the dataset. Correlations with R>0.33 and p<0.0001 were considered significant.

#### Comparison of correlations between male and female mice

R values of all significant correlations were compared with a t-test after Fisher z-transformation. Difference in dependence between parameters between sexes (slope of regression line) was tested by fitting a linear regression model to correlated parameters and assessing significance of interaction between sex and predictor variable. P-values were adjusted for multiple testing with Bonferroni correction to control the family wise error rate to 5%.

#### Immunological structure analysis

Starting from parameters that were affected in a mutant strain, a list was compiled of other parameters with which the affected parameter was ordinarily correlated. The wt C57BL/6N data were then used to predict a value for the correlated parameter in the mutant strain by sex-specific linear regression. If ≥60% of mice for a given mutant strain had predicted values outside the 95% confidence interval based on the wt distribution, the mutant line was defined as breaking this specific parameter correlation. Note that a single perturbed parameter could contribute more than one correlation to the dataset. For the purpose of this analysis, it was assumed that all correlations were equally likely to be broken. To assess if breaking of correlations was correlated with other characteristics, such as a non-immune phenotype or a challenge phenotype, a multivariate Probit regression analysis was undertaken. The same analysis was also used to control for unequal numbers of correlations per gene, allowing for dependencies of correlations within a gene by cluster-robust standard errors and for separate contributions of all characteristics tested (p<0.001 for non-immune controlling for challenge, p<0.001 for challenge controlling for non-immune). The analysis was carried out using R, RStudio and R packages ggplot2, data.table, dplyr and igraph, org.Mm.eg.db and org.Hs.eg.db.

### GnomAD, OMIM and GWAS analyses

If homo-and heterozygotes were analysed from the same strain, only homozygotes were included in this analysis. Human orthologues were based on JAX definition (http://www.informatics.jax.org, accessed 05.05.2016) and only 1:1 human-mouse orthologues were used. GnomAD scores were extracted from release 2 (Hail Table (gs://gnomad-public/release/2.1/ht/constraint/constraint.ht)^47^.

OMIM annotations were obtained on 31 May 2017, with immune-related OMIM-listed genes considered as those implicated in phenotypes of immunodeficiency, recurrent infections, autoimmunity.

GWAS associations were obtained from the full NHGRI-EBI GWAS catalog database 78 on September 10, 2017 (file gwas_catalog_v1.0.1-associations_e89_r2017-07-31.tsv). In GWAS annotation, “immune-related” genes were considered as those mapping to susceptibility to autoimmune disease (systemic lupus erythematosus, psoriasis, rheumatoid arthritis, Sjögren syndrome, primary biliary cholangitis, ulcerative colitis, inflammatory bowel disease, Crohn’s disease, multiple sclerosis, type 1 diabetes, Graves’ disease, late-onset myasthenia gravis); immune response to virus (measured by secreted TNF-alpha); and functional units of gut microbiota.

### Data Availability

The datasets generated and analysed during the current study are available on www.immunophenotype.org and www.mousephenotype.org. Raw FCS flowcytometric files are available from FlowRepository (flowrepository.org). Image files are available from the corresponding author upon request. Code used for initial hit calling and preprocessed per-mouse data for flow cytometry, ear epidermis and DSS assays are available from <https://github.com/AnnaLorenc/3i_heatmapping>.

The following figures have associated raw data which is available from the above-mentioned sources: 1B-D, 2A-C, 4B-C, 5B-E, 6A-D, 7B-F, S1B-H, S2, S3A-D.

All mouse lines analysed in this work are available from repositories linked to the International Mouse Phenotyping Consortium (www.mousephenotype.org) or from WTSI. Cell lines are available upon request.

The PhenStat R package used for Influenza analysis is available on Bioconductor (www.bioconductor.org). The ImageJ macro and the Python code used to score ANA positivity, the Definiens Developer code to assess the ear epidermis images and the R code used to assess the false positive rate are available on request.

## Supplementary Tables 1-5

**Supplementary Table 1.**
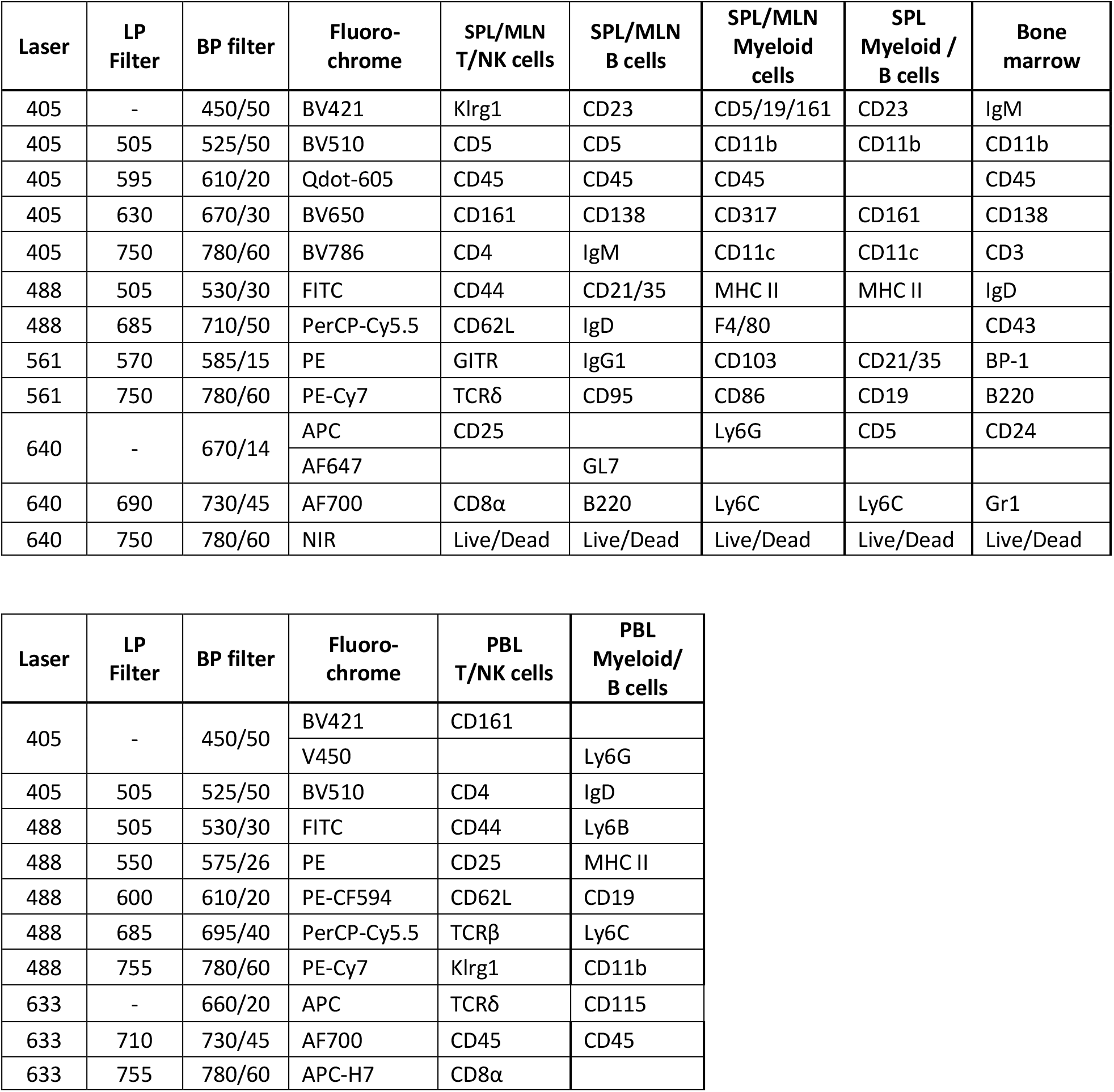
Markers and fluorochromes used to identify immune cell populations by flow cytometry

| Laser | LP Filter | BP filter | Fluoro-chrome | SPL/MLN T/NK cells | SPL/MLN B cells | SPL/MLN Myeloid cells | SPL Myeloid / B cells | Bone marrow |
| --- | --- | --- | --- | --- | --- | --- | --- | --- |
| 405 | - | 450/50 | BV421 | Klrg1 | CD23 | CD5/19/161 | CD23 | IgM |
| 405 | 505 | 525/50 | BV510 | CD5 | CD5 | CD11b | CD11b | CD11b |
| 405 | 595 | 610/20 | Qdot-605 | CD45 | CD45 | CD45 |  | CD45 |
| 405 | 630 | 670/30 | BV650 | CD161 | CD138 | CD317 | CD161 | CD138 |
| 405 | 750 | 780/60 | BV786 | CD4 | IgM | CD11c | CD11c | CD3 |
| 488 | 505 | 530/30 | FITC | CD44 | CD21/35 | MHC II | MHC II | IgD |
| 488 | 685 | 710/50 | PerCP-Cy5.5 | CD62L | IgD | F4/80 |  | CD43 |
| 561 | 570 | 585/15 | PE | GITR | IgG1 | CD103 | CD21/35 | BP-1 |
| 561 | 750 | 780/60 | PE-Cy7 | TCRδ | CD95 | CD86 | CD19 | B220 |
| 640 | - | 670/14 | APC | CD25 |  | Ly6G | CD5 | CD24 |
|  |  |  | AF647 |  | GL7 |  |  |  |
| 640 | 690 | 730/45 | AF700 | CD8α | B220 | Ly6C | Ly6C | Gr1 |
| 640 | 750 | 780/60 | NIR | Live/Dead | Live/Dead | Live/Dead | Live/Dead | Live/Dead |

| Laser | LP Filter | BP filter | Fluoro-chrome | PBL T/NK cells | PBL Myeloid/ B cells |
| --- | --- | --- | --- | --- | --- |
| 405 | - | 450/50 | BV421 | CD161 |  |
|  |  |  | V450 |  | Ly6G |
| 405 | 505 | 525/50 | BV510 | CD4 | IgD |
| 488 | 505 | 530/30 | FITC | CD44 | Ly6B |
| 488 | 550 | 575/26 | PE | CD25 | MHC II |
| 488 | 600 | 610/20 | PE-CF594 | CD62L | CD19 |
| 488 | 685 | 695/40 | PerCP-Cy5.5 | TCRβ | Ly6C |
| 488 | 755 | 780/60 | PE-Cy7 | Klrg1 | CD11b |
| 633 | - | 660/20 | APC | TCRδ | CD115 |
| 633 | 710 | 730/45 | AF700 | CD45 | CD45 |
| 633 | 755 | 780/60 | APC-H7 | CD8α | |

**Supplementary Table 2:** Parameters of flow cytometry and ear epidermis assays

| Parameter | Cell markers |
| --- | --- |
| ***Spleen and mesenteric lymph node*** |  |
| Total $\gamma\delta$ T cells | TCRδ+ |
| CD5+ $\gamma\delta$ T cells | TCRδ+ CD5+ |
| Effector $\gamma\delta$ T cells | TCRδ+ CD62L- CD44+ |
| Resting $\gamma\delta$ T cells | TCRδ+ CD62L+ |
| KLRG1+ $\gamma\delta$ T cells | TCRδ+ KLRG1+ |
| Total $\alpha\beta$ T cells | TCRδ- CD5+ CD4+ or CD8+ |
| Total CD4+ T cells | TCRδ- CD5+ CD4+ |
| CD4+ T helper cells | TCRδ- CD5+ CD4+ CD25- GITR- |
| Effector CD4+ T helper cells | TCRδ- CD5+ CD4+ CD25- GITR- CD62L- CD44+ |
| Resting CD4+ T helper cells | TCRδ- CD5+ CD4+ CD25- GITR- CD62L+ |
| KLRG1+ CD4+ T helper cells | TCRδ- CD5+ CD4+ CD25- GITR- KLRG1+ |
| Total Treg cells | TCRδ- CD5+ CD4+ CD25+ GITR+ |
| Effector Treg cells | TCRδ- CD5+ CD4+ CD25+ GITR+ CD62L- CD44+ |
| Resting Treg cells | TCRδ- CD5+ CD4+ CD25+ GITR+ CD62L+ |
| KLRG1+ Treg cells | TCRδ- CD5+ CD4+ CD25+ GITR+ KLRG1+ |
| Total CD8+ T cells | TCRδ- CD5+ CD8+ |
| Effector CD8+ T cells | TCRδ- CD5+ CD8+ CD62L- CD44+ |
| Resting CD8+ T cells | TCRδ- CD5+ CD8+ CD62L+ CD44+ |
| Naive CD8+ T cells | TCRδ- CD5+ CD8+ CD62L+ CD44- |
| KLRG1+ CD8+ T cells | TCRδ- CD5+ CD8+ KLRG1+ |
| Total NKT cells | TCRδ- CD5+ CD161+ |
| Total CD4- NKT cells | TCRδ- CD5+ CD161+ CD4- |
| Effector CD4- NKT cells | TCRδ- CD5+ CD161+ CD4- CD62L- |
| Resting CD4- NKT cells | TCRδ- CD5+ CD161+ CD4- CD62L+ |
| KLRG1+ CD4- NKT cells | TCRδ- CD5+ CD161+ CD4- KLRG1+ |
| Total CD4+ NKT cells | TCRδ- CD5+ CD161+ CD4+ |
| Effector CD4+ NKT cells | TCRδ- CD5+ CD161+ CD4+ CD62L- |
| Resting CD4+ NKT cells | TCRδ- CD5+ CD161+ CD4+ CD62L+ |
| KLRG1+ CD4+ NKT cells | TCRδ- CD5+ CD161+ CD4+ KLRG1+ |
| Total NK cells | TCRδ- CD5- CD161+ |
| Effector NK cells | TCRδ- CD5- CD161+ CD62L- |
| Resting NK cells | TCRδ- CD5- CD161+ CD62L+ |
| KLRG1+ NK cells | TCRδ- CD5- CD161+ KLRG1+ |
| Plasma cells | CD138+ |
| B1a cells | CD138- B220+ CD5+ |
| B2 cells | CD138- B220+ CD5- |
| Follicular B cells | CD138- B220+ CD5- GL7- CD95- CD21/35<sup>int</sup><br>IgM<sup>int</sup> |
| Memory B cells | CD138- B220+ CD5- GL7- CD95- IgG+ |
| Macrophages | F4/80+ CD11b<sup>int</sup> |
| Granulocytes | F4/80- Ly6G<sup>high</sup> CD11b+ |
| Eosinophils | F4/80- Ly6G<sup>int</sup> CD11b+ SSC<sup>high</sup> |
| Monocytes | F4/80- Ly6G- Ly6C<sup>high</sup> CD11b+ |
| Plasmacytoid DC | F4/80- Ly6G- Ly6C<sup>low/int</sup> CD317+ |
| Conventional DC | F4/80- Ly6G- Ly6C<sup>low/int</sup> CD317- CD11c+ MHCII+ |
| Conventional CD11b<sup>high</sup> DC | F4/80- Ly6G- Ly6C<sup>low/int</sup> CD317- CD11c+ MHCII+<br>CD11b<sup>high</sup> CD86<sup>low</sup> |
| Conventional CD11b<sup>low</sup> DC | F4/80- Ly6G- Ly6C<sup>low/int</sup> CD317- CD11c+ MHCII+<br>CD11b<sup>low</sup> CD86<sup>high</sup> |
| CD103+ conventional CD11b<sup>low</sup> DC | F4/80- Ly6G- Ly6C<sup>low/int</sup> CD317- CD11c+ MHCII+<br>CD11b<sup>low</sup> CD86<sup>high</sup> CD103+ |
| ***Spleen only*** |  |
| Immature NK cells | Ly6G- CD161+ CD5- CD11b- Ly6C- |
| Ly6C+ immature NK cells | Ly6G- CD161+ CD5- CD11b- Ly6C+ |
| Mature NK cells | Ly6G- CD161+ CD5- CD11b+ Ly6C- |
| Ly6C+ mature NK cells | Ly6G- CD161+ CD5- CD11b+ Ly6C+ |
| Germinal centre B cells | CD138- B220+ CD5- GL7+ CD95+ |
| Early germinal centre B cells | CD138- B220+ CD5- GL7+ CD95+ IgM+ IgG- |
| Late germinal centre B cells | CD138- B220+ CD5- GL7+ CD95+ IgM- IgG+ |
| Marginal zone precursor B cells | CD138- B220+ CD5- GL7- CD95- CD21/35+<br>CD23- |
| Marginal zone B cells | CD138- B220+ CD5- GL7- CD95- CD21/35+<br>CD23+ |
| Transitional 1 B cells | CD138- B220+ CD5- GL7- CD95- CD21/35-<br>CD23- |
| Transitional 2 B cells | CD138- B220+ CD5- GL7- CD95- CD21/35-<br>CD23+ |
| ***Blood*** |  |
| Total T cells | TCRβ+ or TCRδ+ |
| Total $\alpha\beta$ T cells | TCRβ+ |
| Total CD4+ T cells | TCRβ+ CD4+ |
| Effector CD4+ T cells | TCRβ+ CD4+ CD62L- CD44+ |
| KLRG1+ CD4+ T cells | TCRβ+ CD4+ KLRG1+ |
| Total Treg cells | TCRβ+ CD4+ CD25+ |
| Total CD8+ T cells | TCRβ+ CD8+ |
| Effector CD8+ T cells | TCRβ+ CD8+ CD62L- CD44+ |
| KLRG1+ CD8+ T cells | TCRβ+ CD8+ KLRG1+ |
| Total $\gamma\delta$ T cells | TCRδ+ |
| Total NKT cells | TCRβ+ CD161+ |
| Total NK cells | TCRβ- CD161+ |
| KLRG1+ NK cells | TCRβ- CD161+ KLRG1+ |
| Total B cells | CD19+ |
| IgD+ mature B cells | CD19+ IgD+ |
| Monocytes | CD19- Ly6G- CD11b+ |
| Ly6C+ monocytes | CD19- Ly6G- CD11b+ Ly6C+ |
| Ly6C- monocytes | CD19- Ly6G- CD11b+ Ly6C- |
| Neutrophils | CD19- Ly6G+ Ly6B+ |
| Eosinophils | CD19- Ly6B- Ly6G- SSC<sup>high</sup> CD11b+ |
| ***Bone marrow*** |  |
| Granulocytes | GR+ CD43+ |
| T cells | GR- CD3+ |
| Plasma cells | GR- CD3- CD138+ |
| Myeloid cells | GR- CD3- CD138- CD11b+ |
| Total B cell precursors | GR- CD3- CD138- B220+ |
| Pre-pro B cells (Hardy fraction A) | GR- CD3- CD138- B220+ CD43+ BP1- CD24- |
| Pro-B cells (Hardy fraction B) | GR- CD3- CD138- B220+ CD43+ BP1- CD24+ |
| Pro-B cells (Hardy fraction C) | GR- CD3- CD138- B220+ CD43+ BP1+ CD24+ |
| Pre-B cells (Hardy fraction D) | GR- CD3- CD138- B220+ CD43- IgD- IgM- |
| Immature B cells (Hardy fraction E) | GR- CD3- CD138- B220+ CD43- IgD- IgM+ |
| Mature B cells (Hardy fraction F) | GR- CD3- CD138- B220+ CD43- IgD+ IgM+ |
| ***Ear epidermis*** |  |
| DETC coverage |  |
| Count of DETC Vγ5+ | |
| Count of atypical CD45+ cells |  |
| Count of DETC Vγ5+ contacting LC | |
| Count of DETC Vγ5+ not contacting LC | |
| Count of LC |  |
| Average volume of DETC Vγ5+ | |
| Average volume of atypical CD45+ cells |  |
| Average volume of DETC Vγ5+ contacting LC | |
| Average volume of DETC Vγ5+ not contacting LC | |
| Average volume of LC |  |
| Average roundness of DETC Vγ5+ | |
| Average roundness of atypical CD45+ cells |  |
| Average roundness of DETC Vγ5+ contacting LC | |
| Average roundness of DETC Vγ5+ not contacting LC | |
| Average roundness of LC |  |
| Average number of branches on DETC Vγ5+ | |
| Average number of branches on DETC Vγ5+ contacting LC | |
| Average number of branches on DETC Vγ5+ not contacting LC | |
| Average number of branches on LC |  |
| Average branch length of on DETC Vγ5+ | |
| Average branch length of on DETC Vγ5+ contacting LC | |
| Average branch length of on DETC Vγ5+ not contacting LC | |

**Supplementary Table 3:**
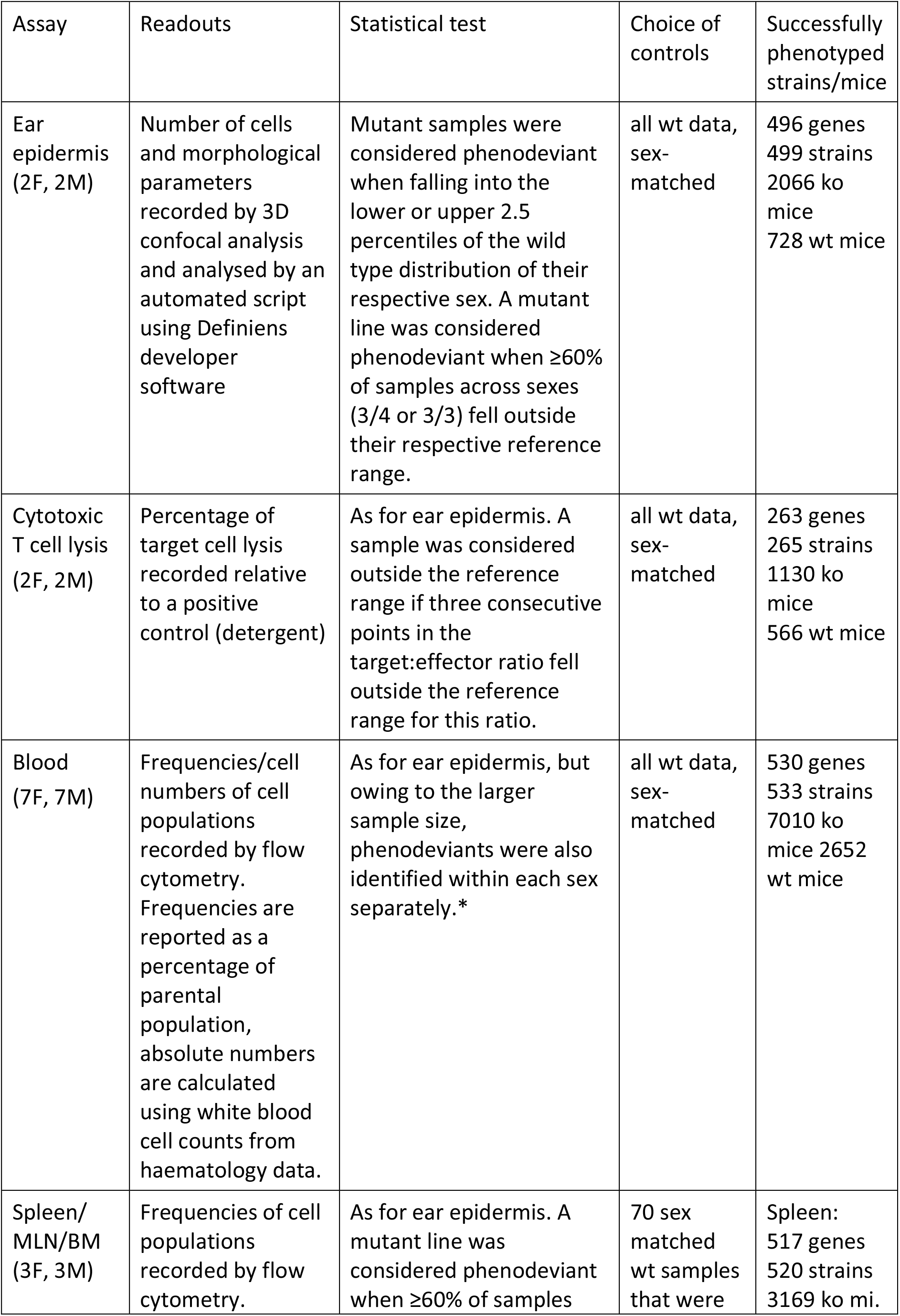
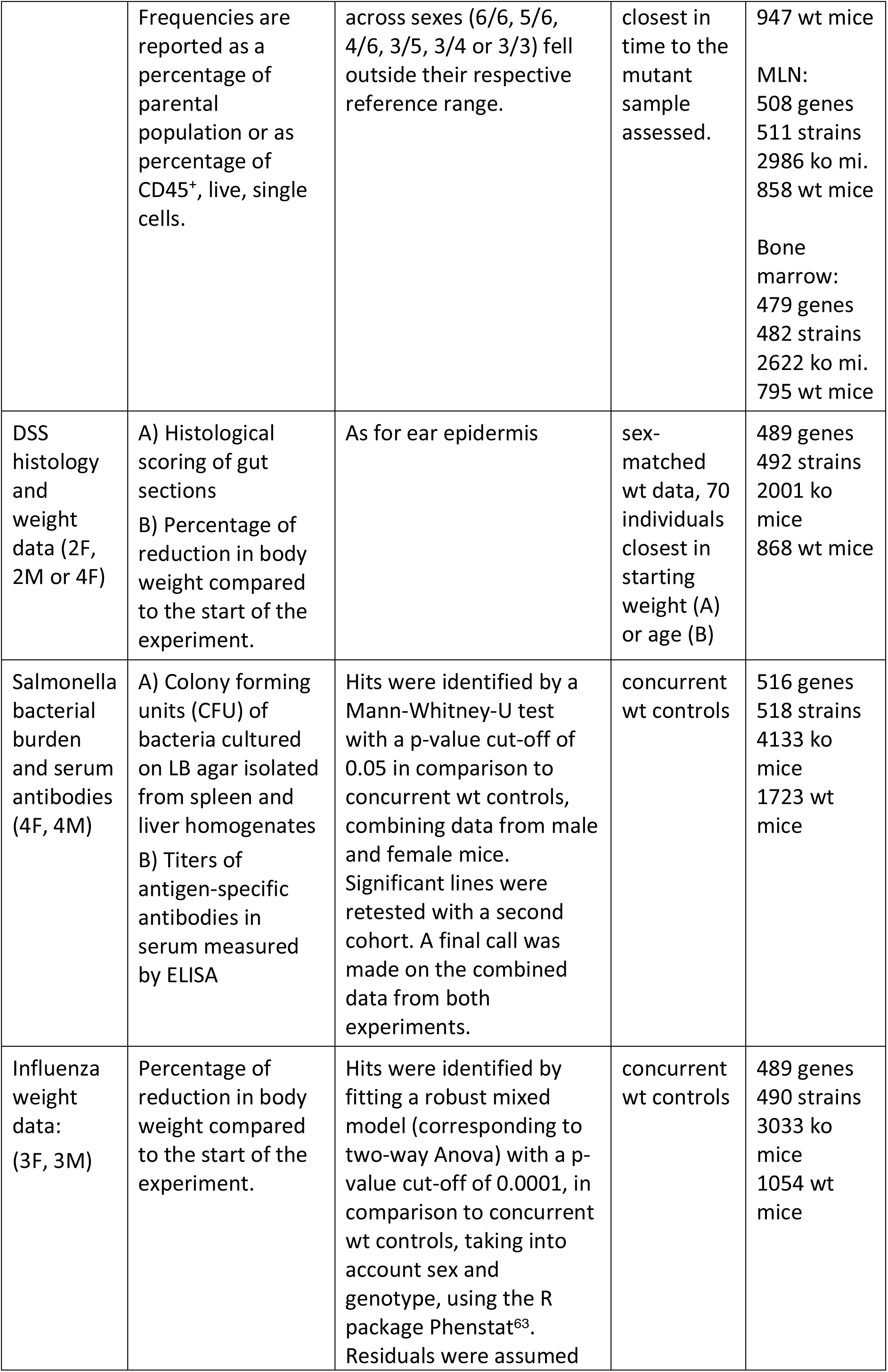
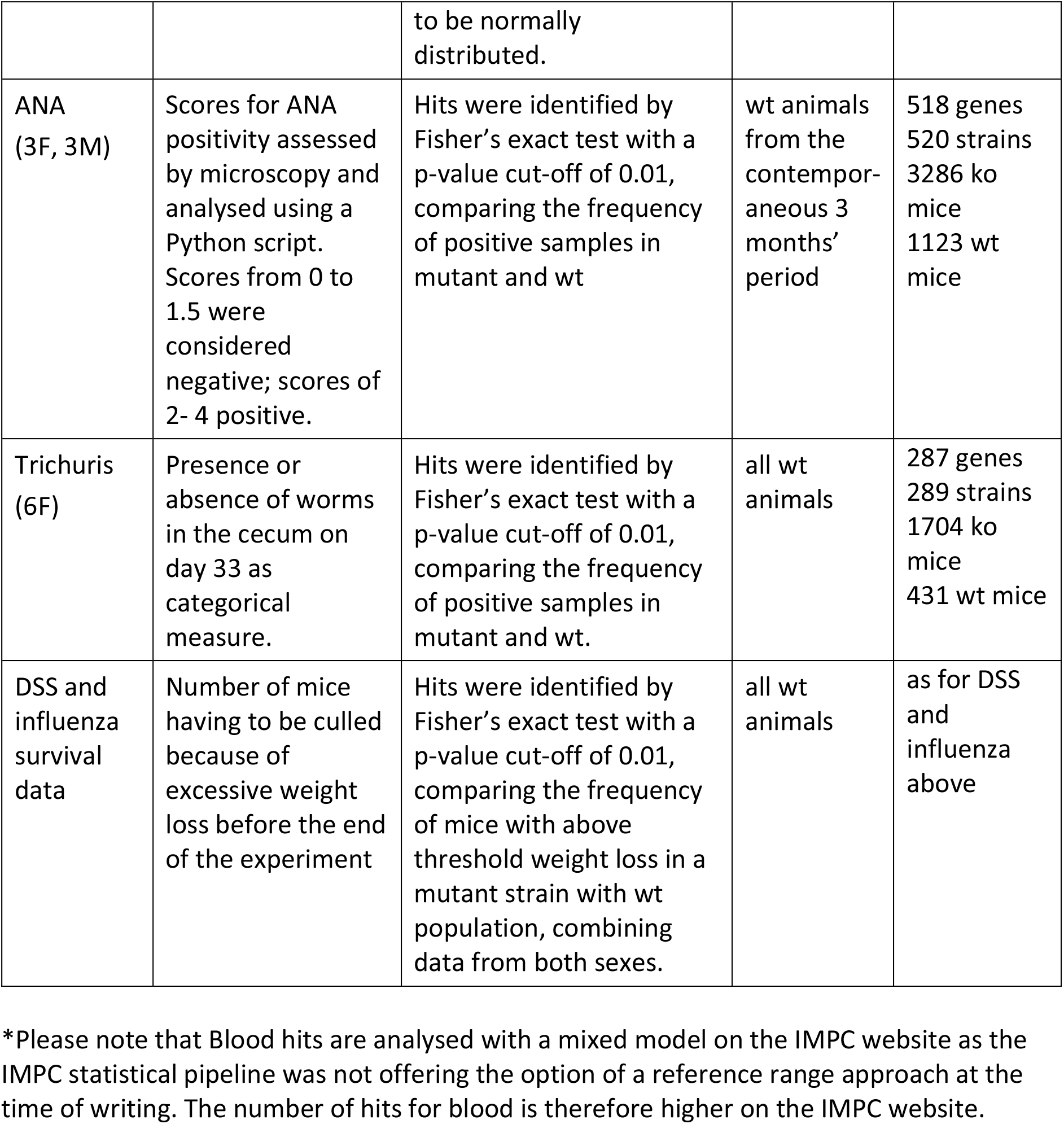
Summary of statistical procedures per assay and readout

| Assay | Readouts | Statistical test | Choice of controls | Successfully phenotyped strains/mice |
| --- | --- | --- | --- | --- |
| Ear epidermis (2F, 2M) | Number of cells and morphological parameters recorded by 3D confocal analysis and analysed by an automated script using Definiens developer software | Mutant samples were considered phenodeviant when falling into the lower or upper 2.5 percentiles of the wild type distribution of their respective sex. A mutant line was considered phenodeviant when $\geq 60\%$ of samples across sexes (3/4 or 3/3) fell outside their respective reference range. | all wt data, sex-matched | 496 genes<br>499 strains<br>2066 ko mice<br>728 wt mice |
| Cytotoxic T cell lysis (2F, 2M) | Percentage of target cell lysis recorded relative to a positive control (detergent) | As for ear epidermis. A sample was considered outside the reference range if three consecutive points in the target:effector ratio fell outside the reference range for this ratio. | all wt data, sex-matched | 263 genes<br>265 strains<br>1130 ko mice<br>566 wt mice |
| Blood (7F, 7M) | Frequencies/cell numbers of cell populations recorded by flow cytometry. Frequencies are reported as a percentage of parental population, absolute numbers are calculated using white blood cell counts from haematology data. | As for ear epidermis, but owing to the larger sample size, phenodeviants were also identified within each sex separately.* | all wt data, sex-matched | 530 genes<br>533 strains<br>7010 ko mice<br>2652 wt mice |
| Spleen/MLN/BM (3F, 3M) | Frequencies of cell populations recorded by flow cytometry. | As for ear epidermis. A mutant line was considered phenodeviant when $\geq 60\%$ of samples | 70 sex matched wt samples that were | Spleen:<br>517 genes<br>520 strains<br>3169 ko mi. |
|  | Frequencies are reported as a percentage of parental population or as percentage of CD45<sup>+</sup>, live, single cells. | across sexes (6/6, 5/6, 4/6, 3/5, 3/4 or 3/3) fell outside their respective reference range. | closest in time to the mutant sample assessed. | 947 wt mice<br>MLN:<br>508 genes<br>511 strains<br>2986 ko mi.<br>858 wt mice<br>Bone marrow:<br>479 genes<br>482 strains<br>2622 ko mi.<br>795 wt mice |
| DSS histology and weight data (2F, 2M or 4F) | A) Histological scoring of gut sections<br>B) Percentage of reduction in body weight compared to the start of the experiment. | As for ear epidermis | sex-matched wt data, 70 individuals closest in starting weight (A) or age (B) | 489 genes<br>492 strains<br>2001 ko mice<br>868 wt mice |
| Salmonella bacterial burden and serum antibodies (4F, 4M) | A) Colony forming units (CFU) of bacteria cultured on LB agar isolated from spleen and liver homogenates<br>B) Titers of antigen-specific antibodies in serum measured by ELISA | Hits were identified by a Mann-Whitney-U test with a p-value cut-off of 0.05 in comparison to concurrent wt controls, combining data from male and female mice. Significant lines were retested with a second cohort. A final call was made on the combined data from both experiments. | concurrent wt controls | 516 genes<br>518 strains<br>4133 ko mice<br>1723 wt mice |
| Influenza weight data: (3F, 3M) | Percentage of reduction in body weight compared to the start of the experiment. | Hits were identified by fitting a robust mixed model (corresponding to two-way Anova) with a p-value cut-off of 0.0001, in comparison to concurrent wt controls, taking into account sex and genotype, using the R package Phenstat<sup>63</sup>. Residuals were assumed to be normally distributed. | concurrent wt controls | 489 genes<br>490 strains<br>3033 ko mice<br>1054 wt mice |
| ANA<br>(3F, 3M) | Scores for ANA positivity assessed by microscopy and analysed using a Python script. Scores from 0 to 1.5 were considered negative; scores of 2- 4 positive. | Hits were identified by Fisher's exact test with a p-value cut-off of 0.01, comparing the frequency of positive samples in mutant and wt | wt animals from the contemporaneous 3 months' period | 518 genes<br>520 strains<br>3286 ko mice<br>1123 wt mice |
| Trichuris<br>(6F) | Presence or absence of worms in the cecum on day 33 as categorical measure. | Hits were identified by Fisher's exact test with a p-value cut-off of 0.01, comparing the frequency of positive samples in mutant and wt. | all wt animals | 287 genes<br>289 strains<br>1704 ko mice<br>431 wt mice |
| DSS and influenza survival data | Number of mice having to be culled because of excessive weight loss before the end of the experiment | Hits were identified by Fisher's exact test with a p-value cut-off of 0.01, comparing the frequency of mice with above threshold weight loss in a mutant strain with wt population, combining data from both sexes. | all wt animals | as for DSS and influenza above |

\*Please note that Blood hits are analysed with a mixed model on the IMPC website as the IMPC statistical pipeline was not offering the option of a reference range approach at the time of writing. The number of hits for blood is therefore higher on the IMPC website.

**Supplementary Table 4:**
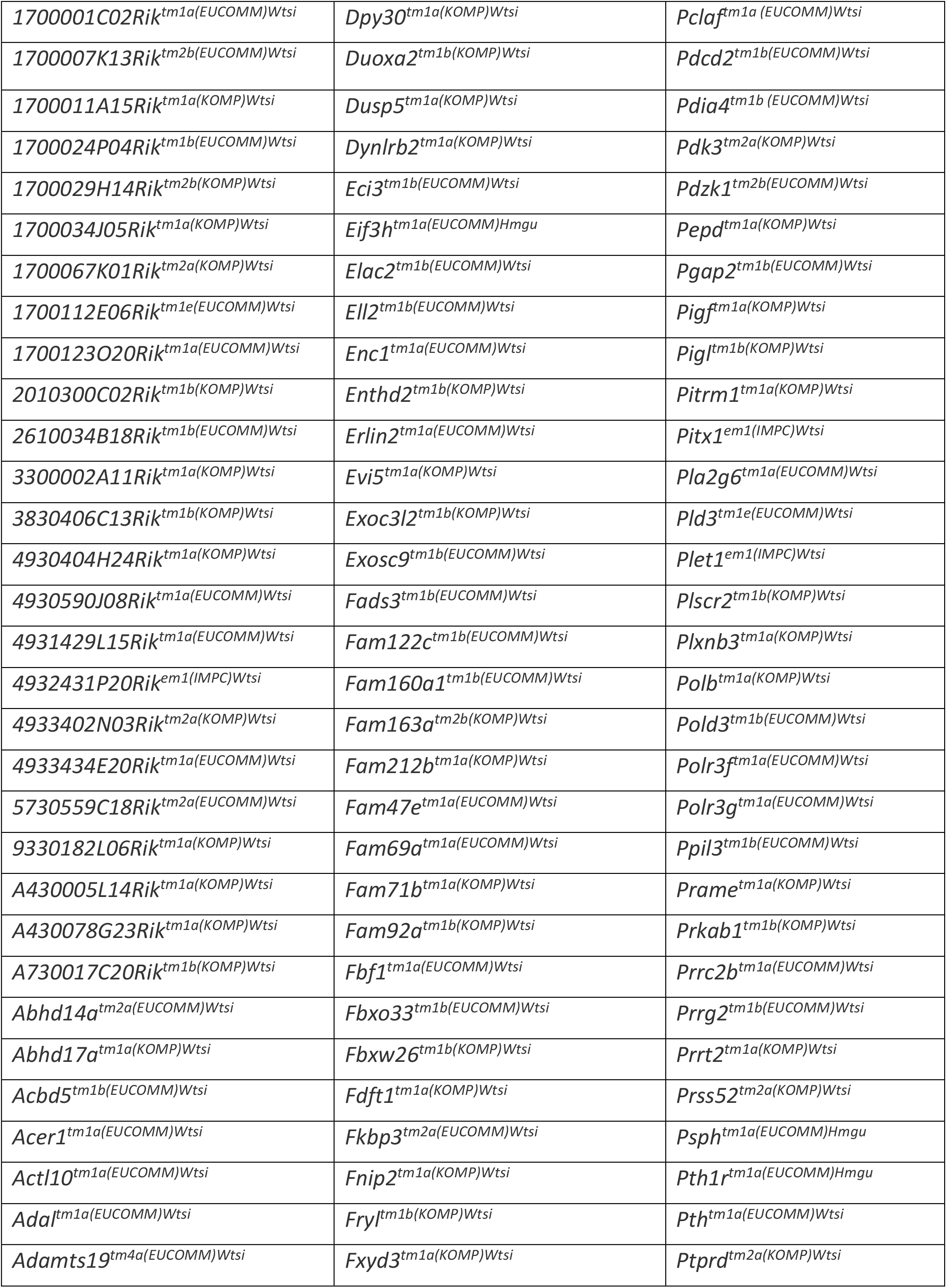
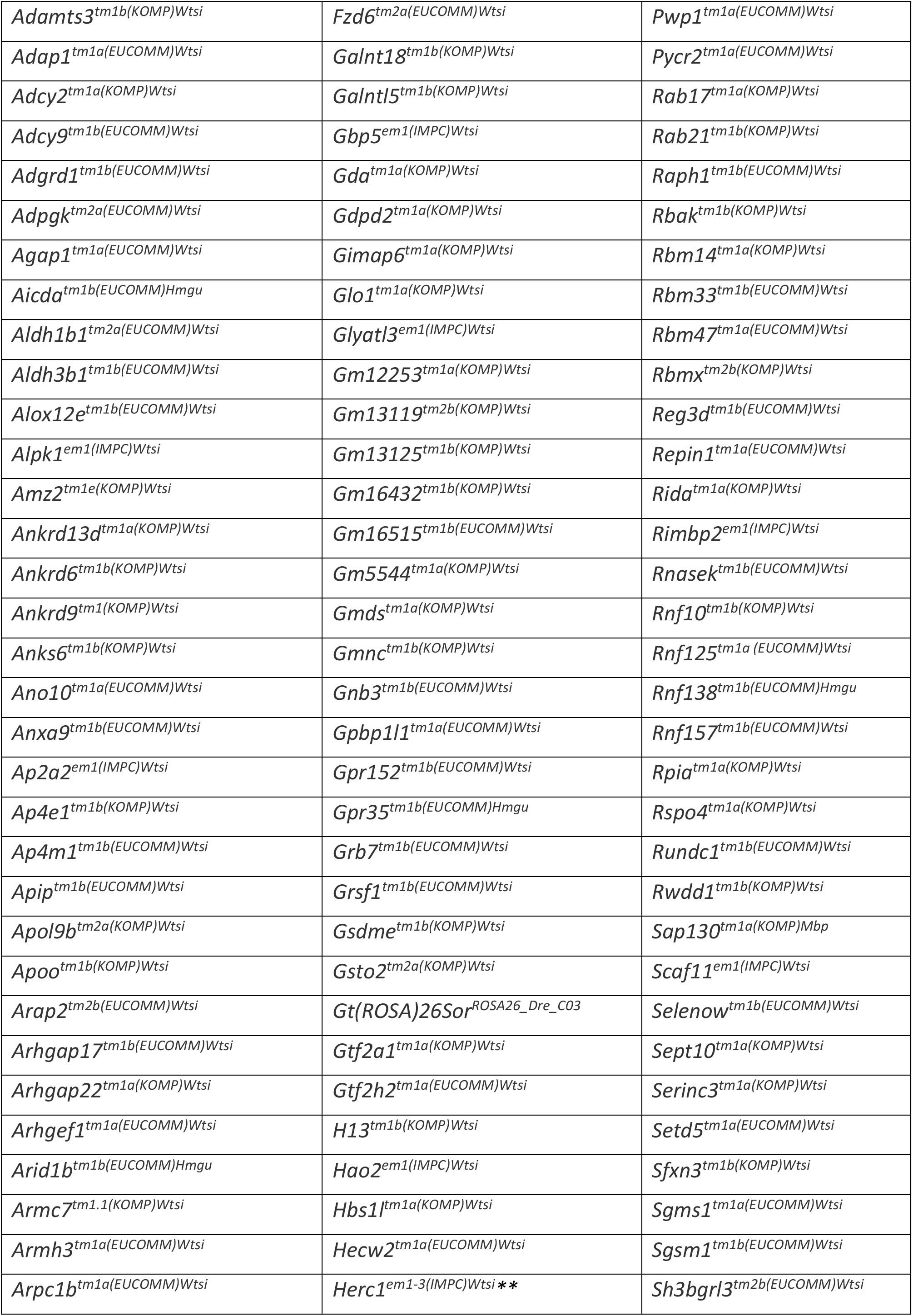
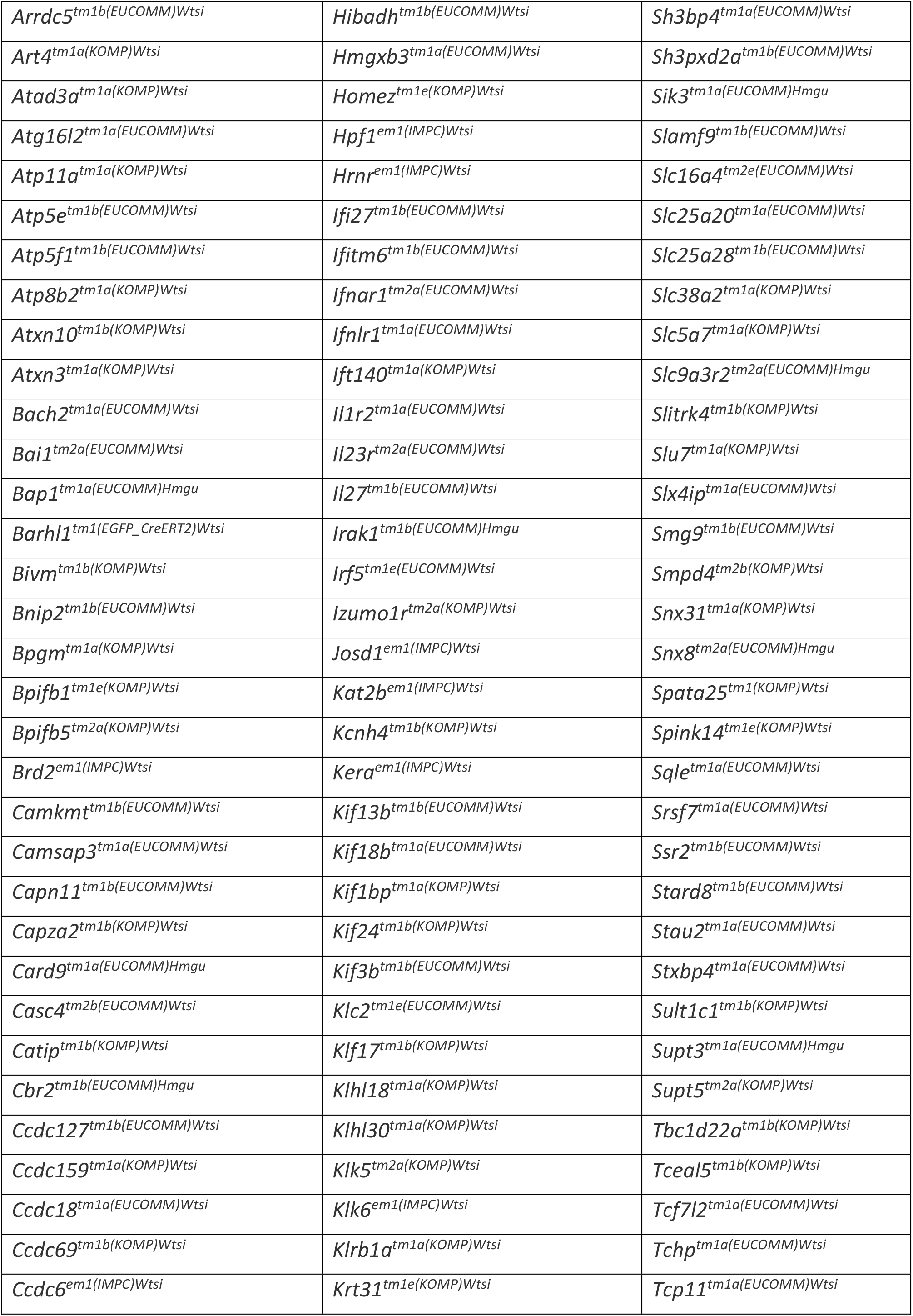
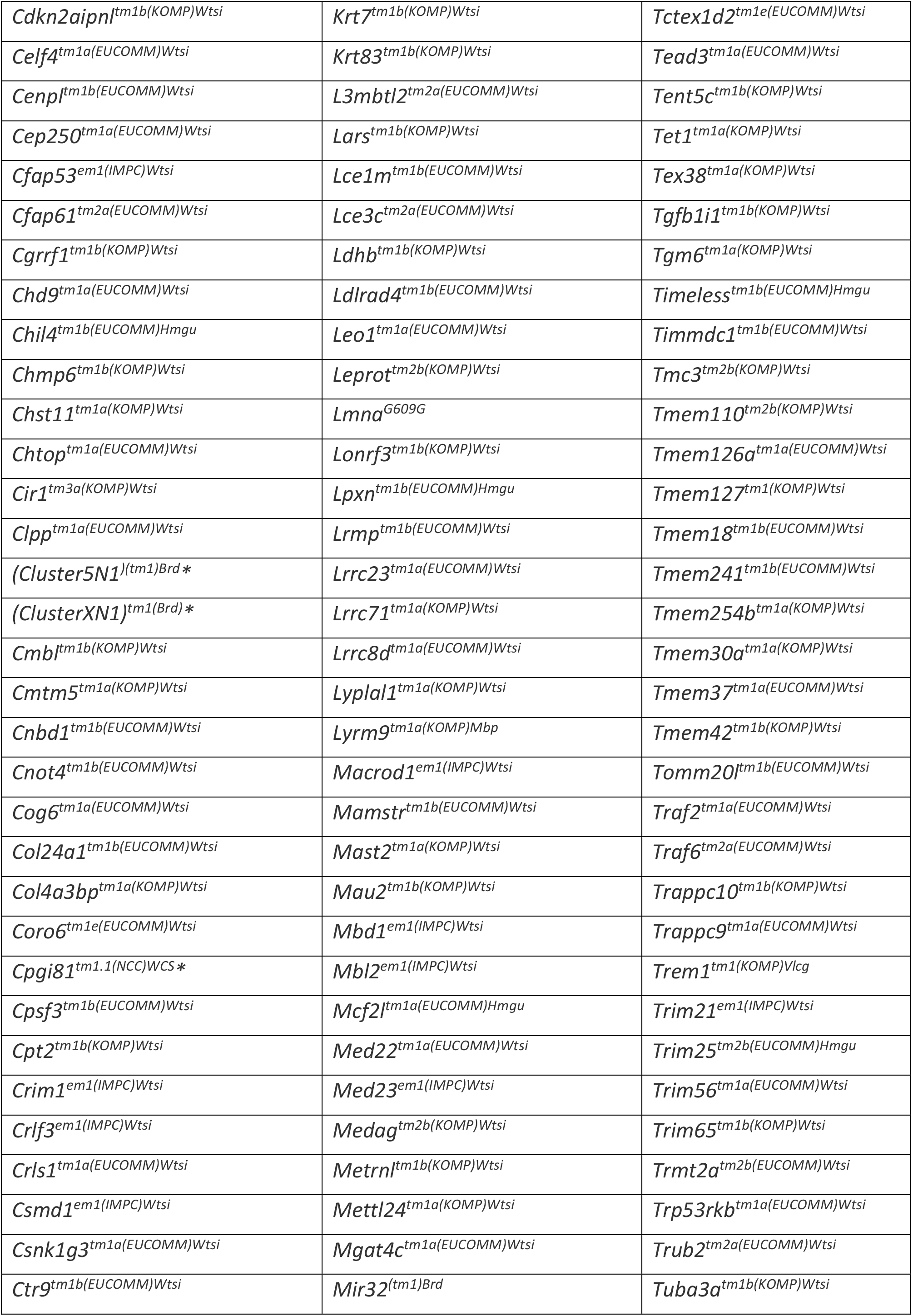
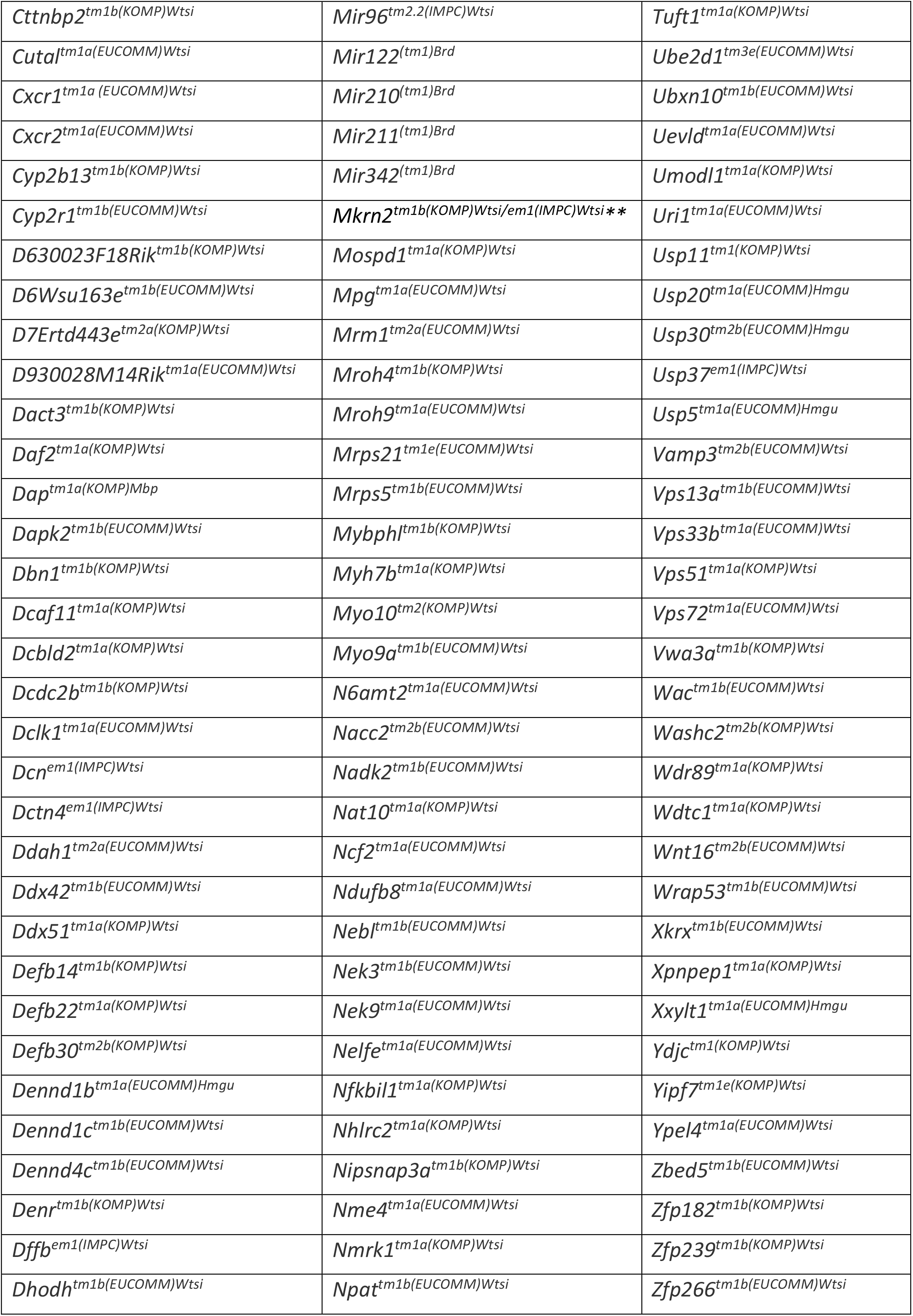
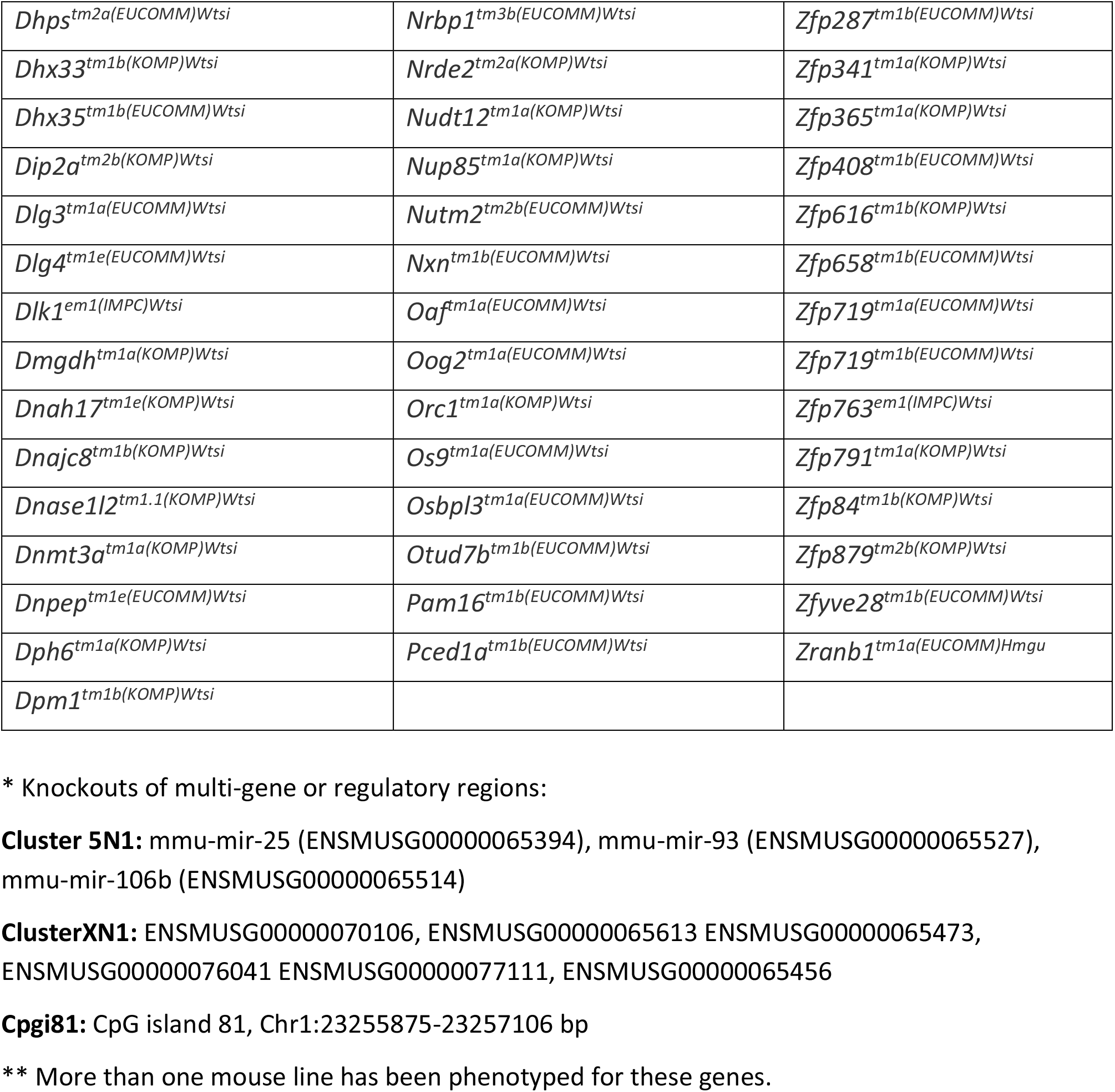
Alleles of knockout lines included in the study

**Supplementary Table 5:**
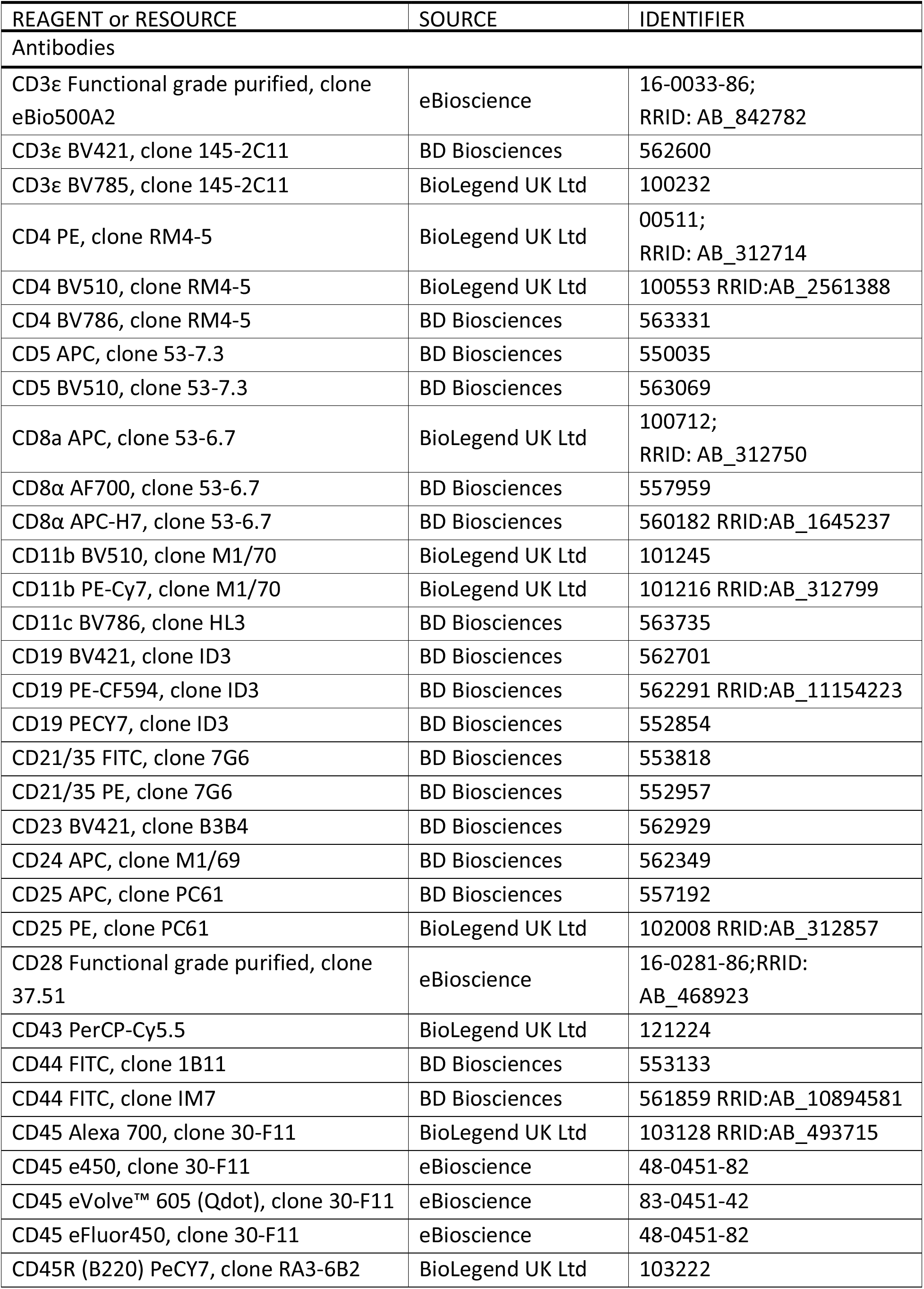
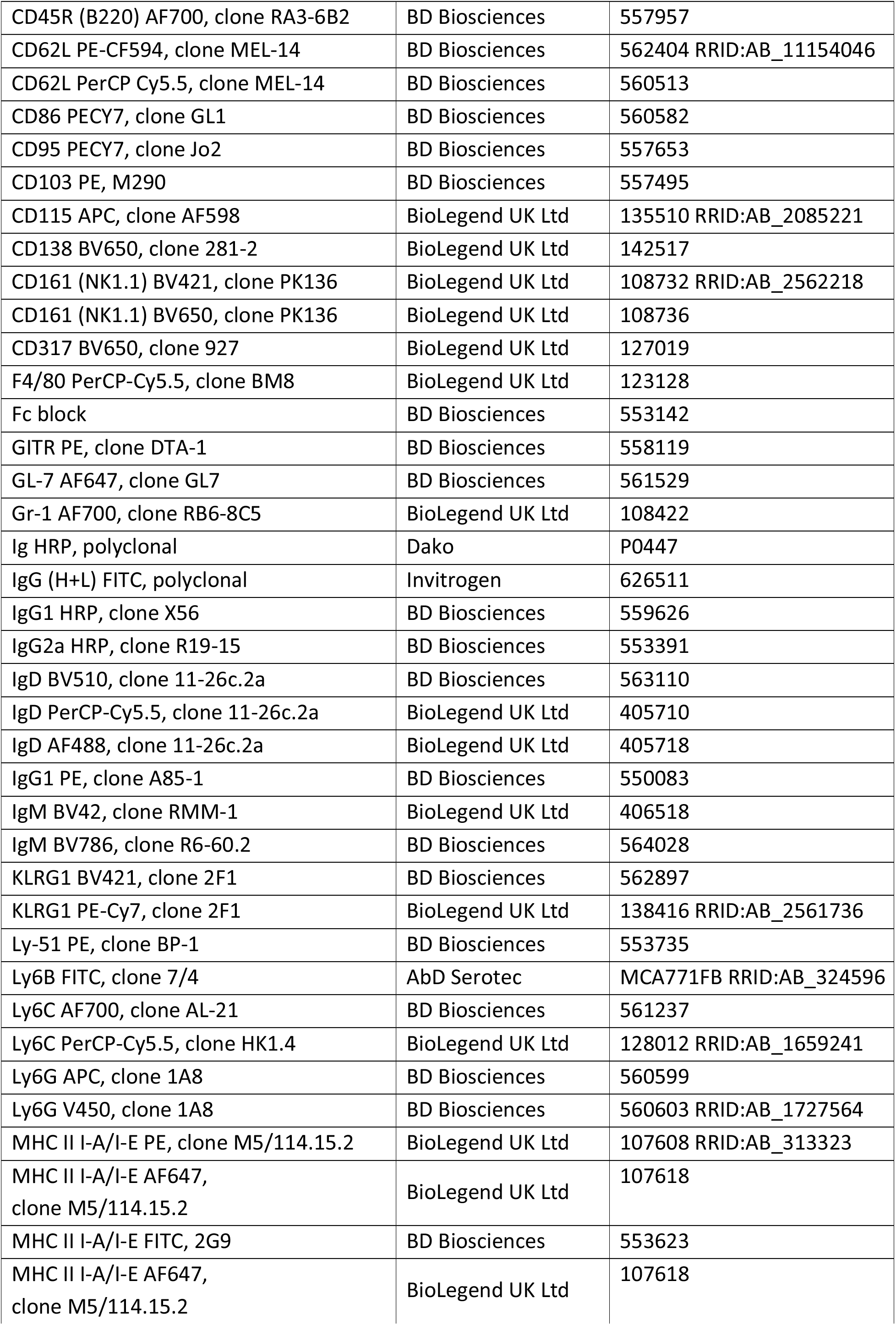
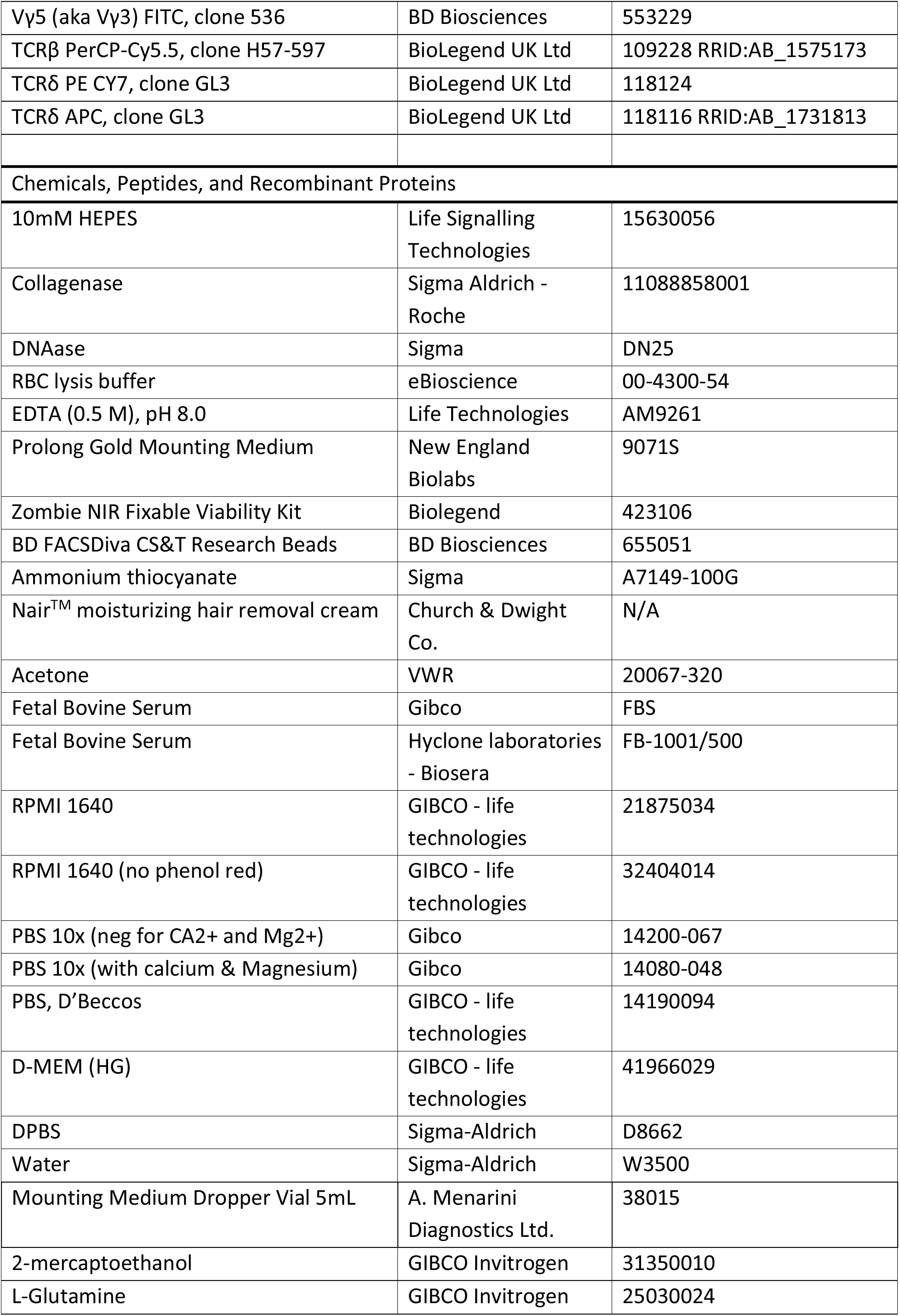
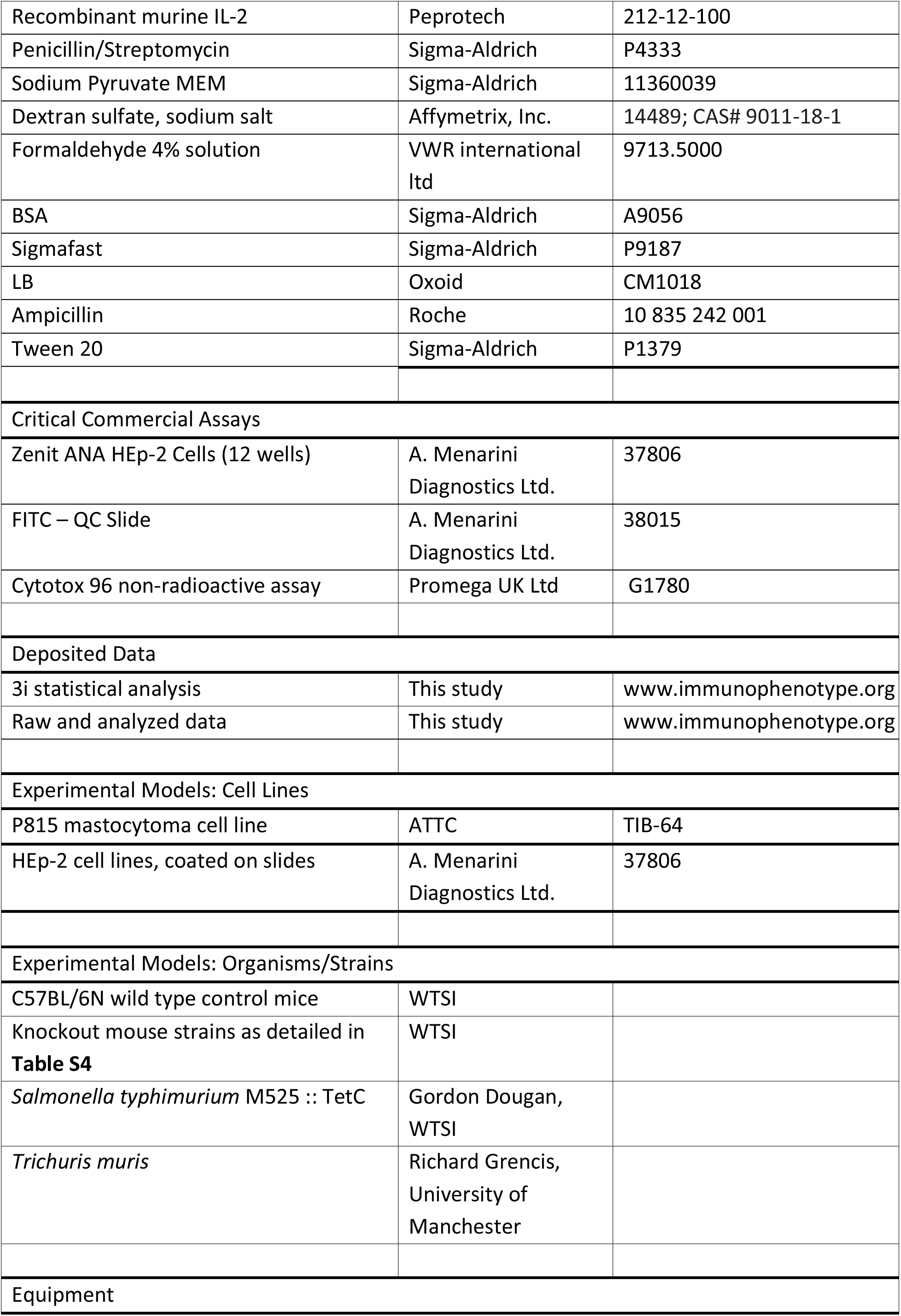
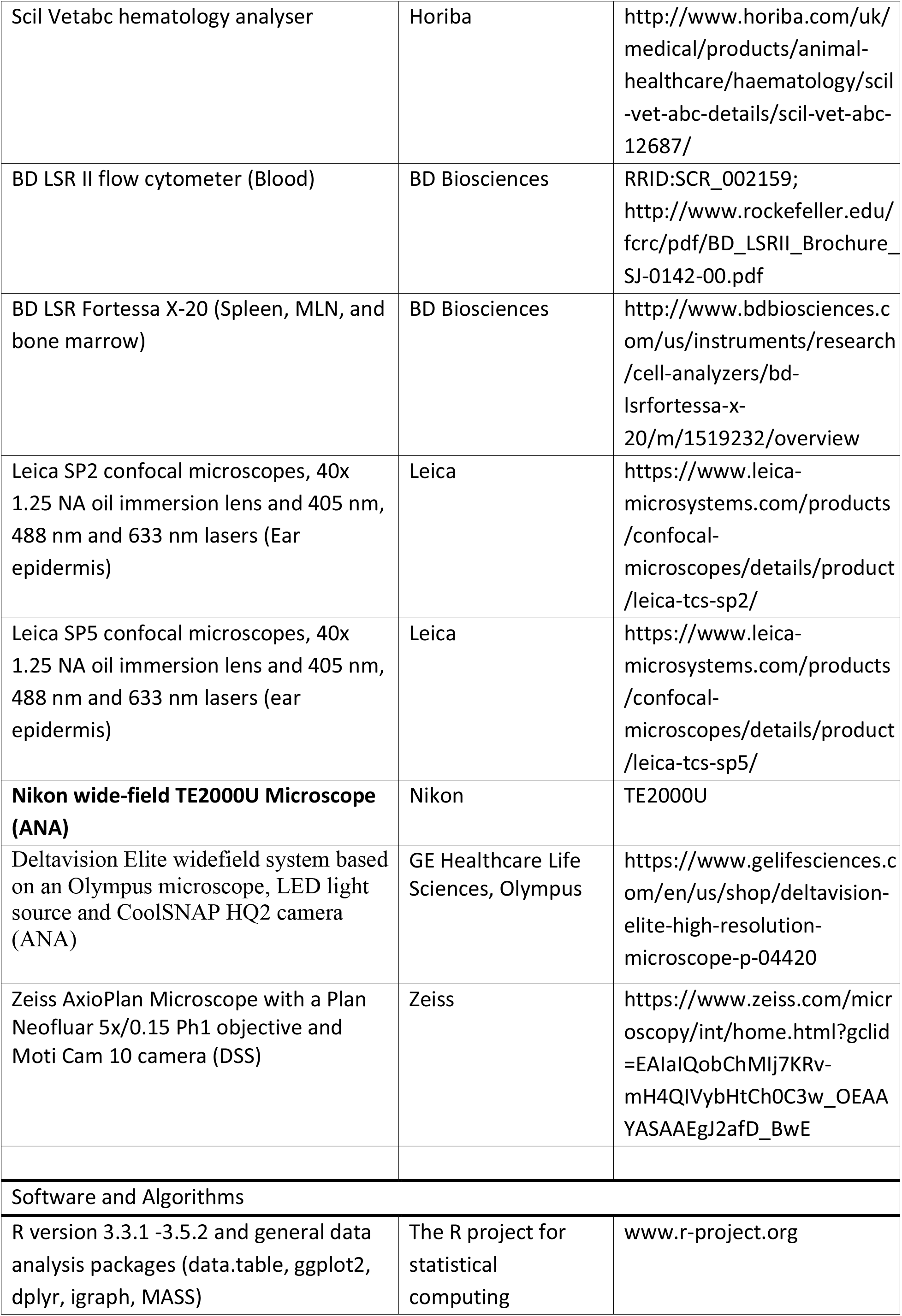
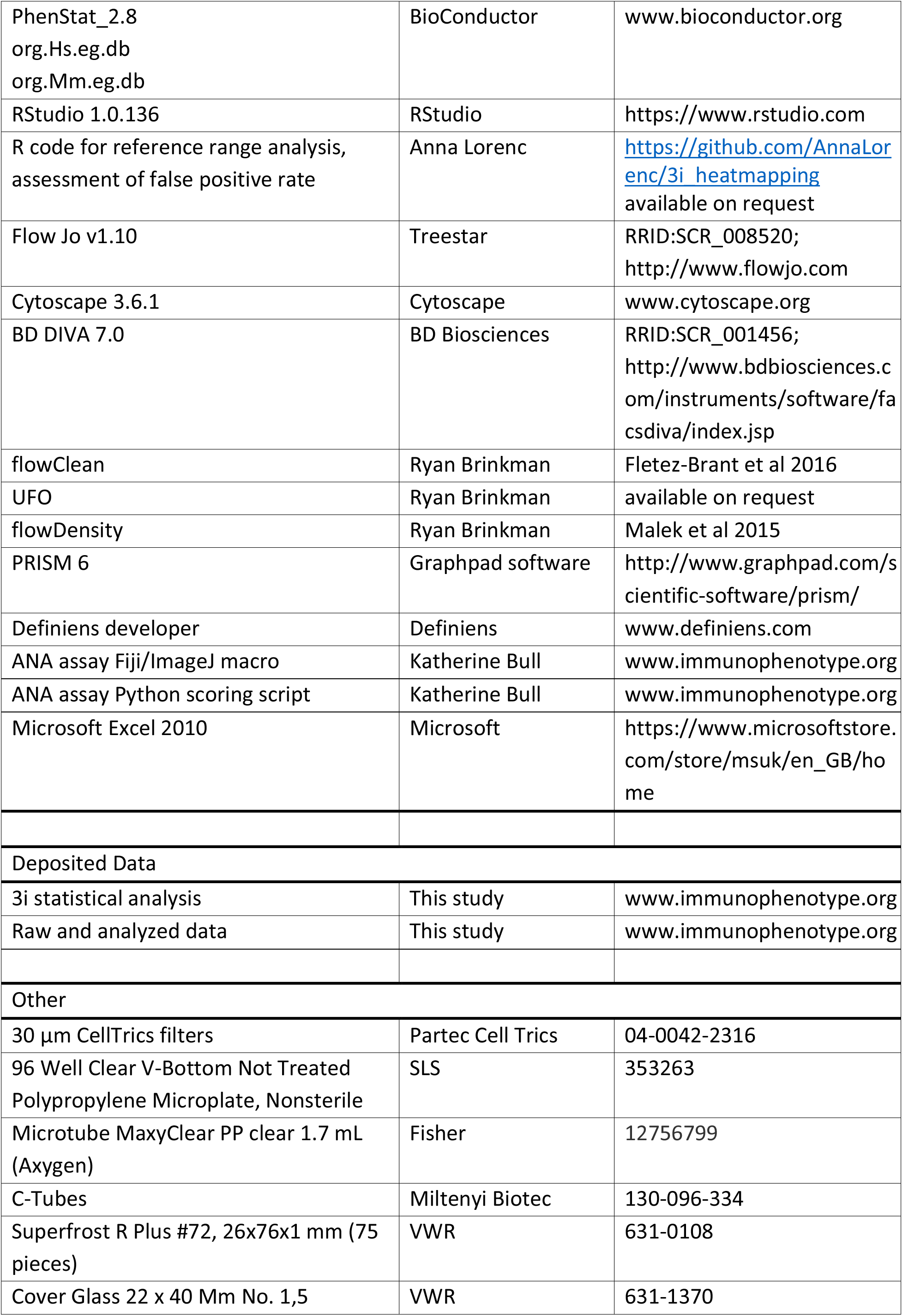
Key resources

**Supplementary Figure 1:**
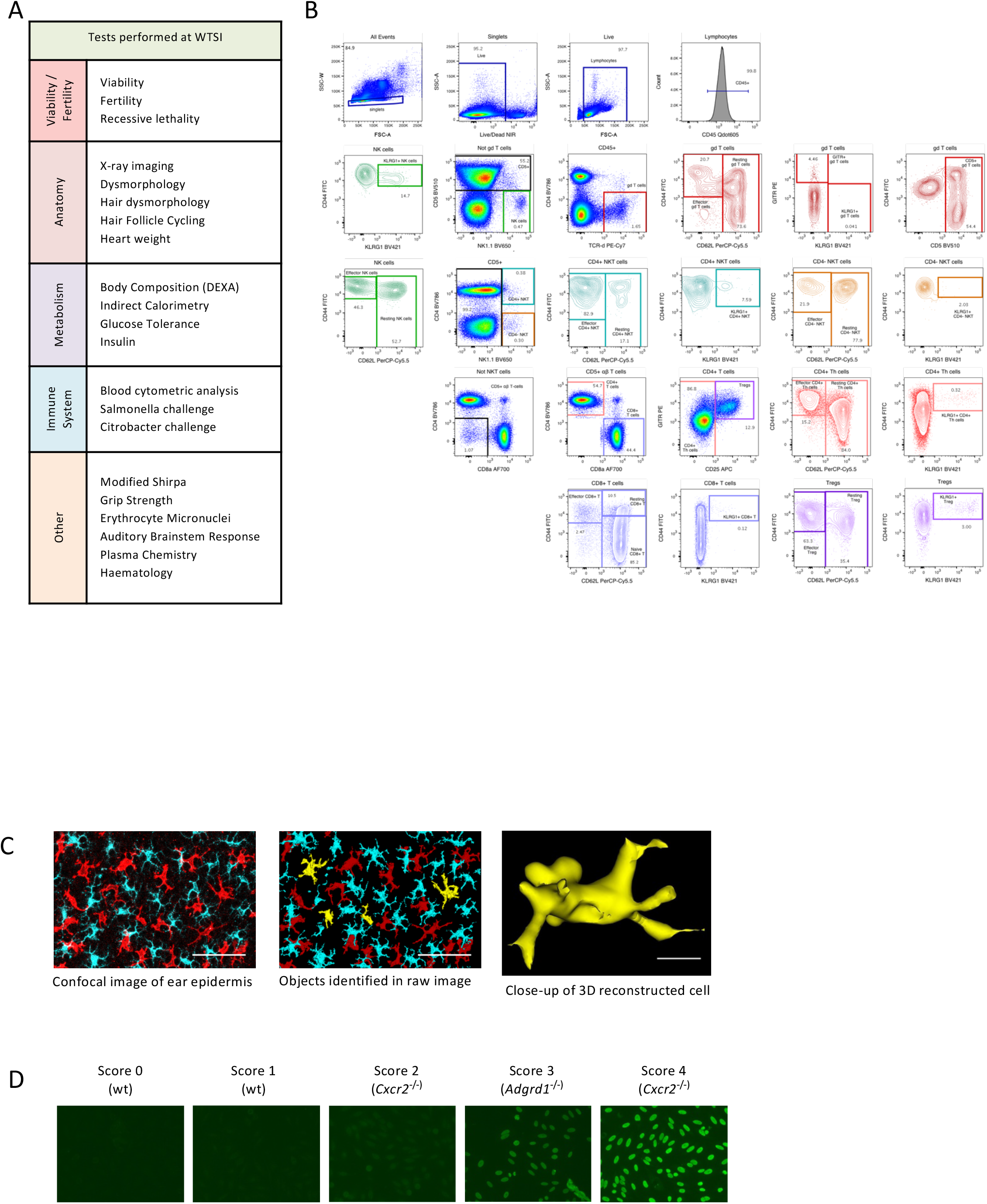
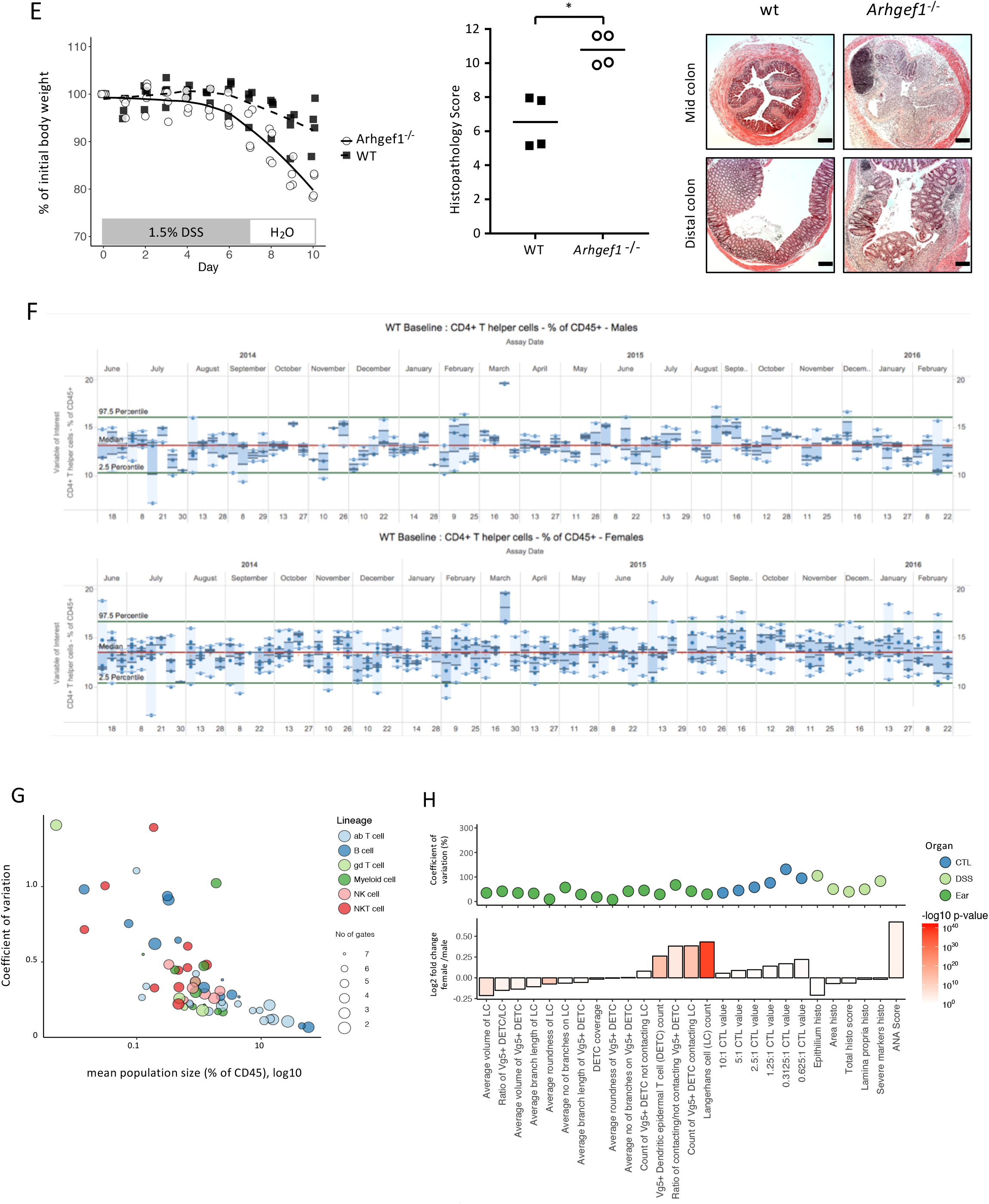
Assay overview. A. Overview of the IMPC tests performed at Wellcome Trust Sanger Institute (WTSI) prior to the substantial expansion by 3i. B. Flow cytometry screen. Representative MLN data illustrate gating strategy for T cells, NKT cells and NK cells. Subsets of key populations are shown in gates of the same colour. Percentages denote percentages of parent population. Panels can be found in **Table S1**. C. Epidermis screen. Left panel: Maximal intensity projection of a z-stack 2-channel overlay. Pseudocolouring scheme was applied to represent Dendritic Epidermal Vγ5^+^ γδ T Cells (DETC) in red and MHC-II+ Langerhans cells (LC) in cyan, scale bar 50 μm. Middle panel: 3D quantitative object-based image analysis (Definiens Developer XD) of DETC that contact LC (red) versus not contacting LC (yellow), and LC (cyan); scale bar: 50μm. Right panel: 3D representation of a DETC rendering in Definiens Developer from which parameters such as dendricity and roundness were quantitated; scale bar: 10μm. D. ANA screen. Antinuclear antibody staining in Hep2 cells using serum from wt mice and different ko lines. Sera were scored on a scale of 0 to 4. The images illustrate that sera from wt or specific mutant strains do not all score equally. Scores of ≤1 were regarded as negative; scores of ≥2 as positive. E. DSS histology screen. Left: Body weight curves of wt and *Arhgef1^-/-^* mice in the same experiment (n=4[2M2F] for wt; n=4[2M2f] for *Arhgef1^-/-^*). Middle: histopathology scores of wt and *Arhgef1^-/-^* mice. Symbols denote the average histopathology score for mid- and distal-colon in each mouse, and horizontal bars represent the median across mice for this group. Right: representative photomicrographs of H&E-stained colonic sections of wt and *Arhgef1^-/-^* mice; scale bar 0.2 mm. F. Longitudinal monitoring of manually analysed flow cytometry data for quality controls (QC), illustrated by CD4^+^ T helper cells. Inner quantiles are shaded dark blue, outer quantiles light blue. Temporal QC are further considered in **Methods**. G. Variation in immune cell subsets (colours denote cell lineages) detected by flow cytometry is generally greater in smaller populations, but does not correlate with number of gates by which populations are defined (denoted by size of circles). Data from 16-week old wt C57BL/6N SPL (n>500). H. Variation and sexual dimorphism in analyses of ear epidermal microscopy; CD8 cytotoxicity; DSS colitis; and ANA. Lower panel: sexual dimorphism of mean values as log2 fold change (middle panel, female/male); colour indicates p value. Upper panel: coefficient of variation; colour indicates assay. Number of wt samples for each of the assays are listed in **Table S3**.

**Supplementary Figure 2:**
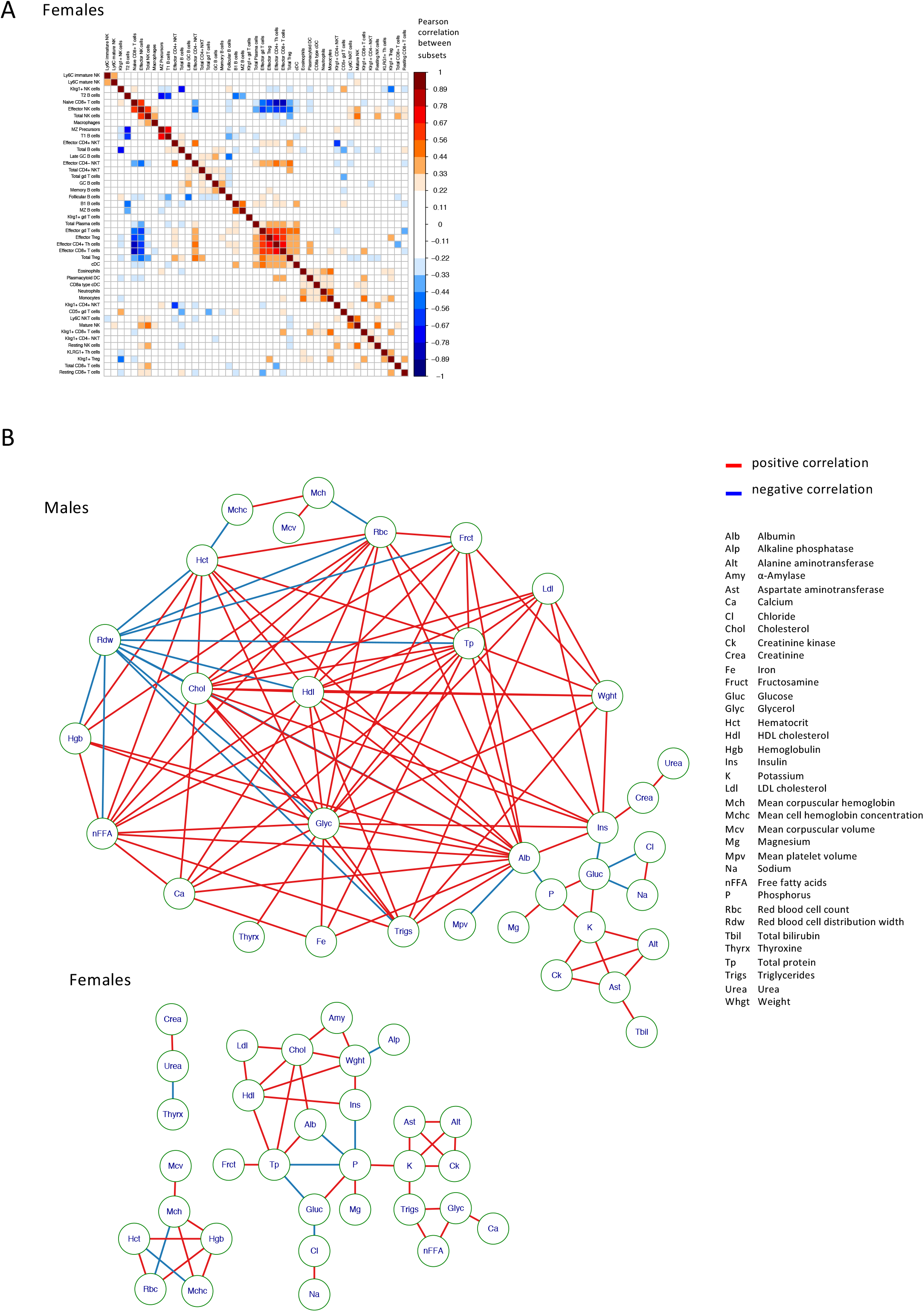
immunological and non-immunological correlations across mice. A. Several SPL cell subsets correlate with each other. Heat map represents Pearson correlations of 46 splenic immune cell subsets with each other in wt females (n>230) as determined by flow cytometry. Dark red fields denote strong positive, dark blue fields strong negative correlations. B. Many non-immune parameters are densely correlated with each other. Correlations of nonimmune parameters (haematological, clinical blood chemistry, and additional data) in males and females. Depicted are correlations with an R-value >0.33 and p<0.001; red lines denote positive correlations; blue lines, negative correlations (n>780 per sex).

**Supplementary Figure 3:**
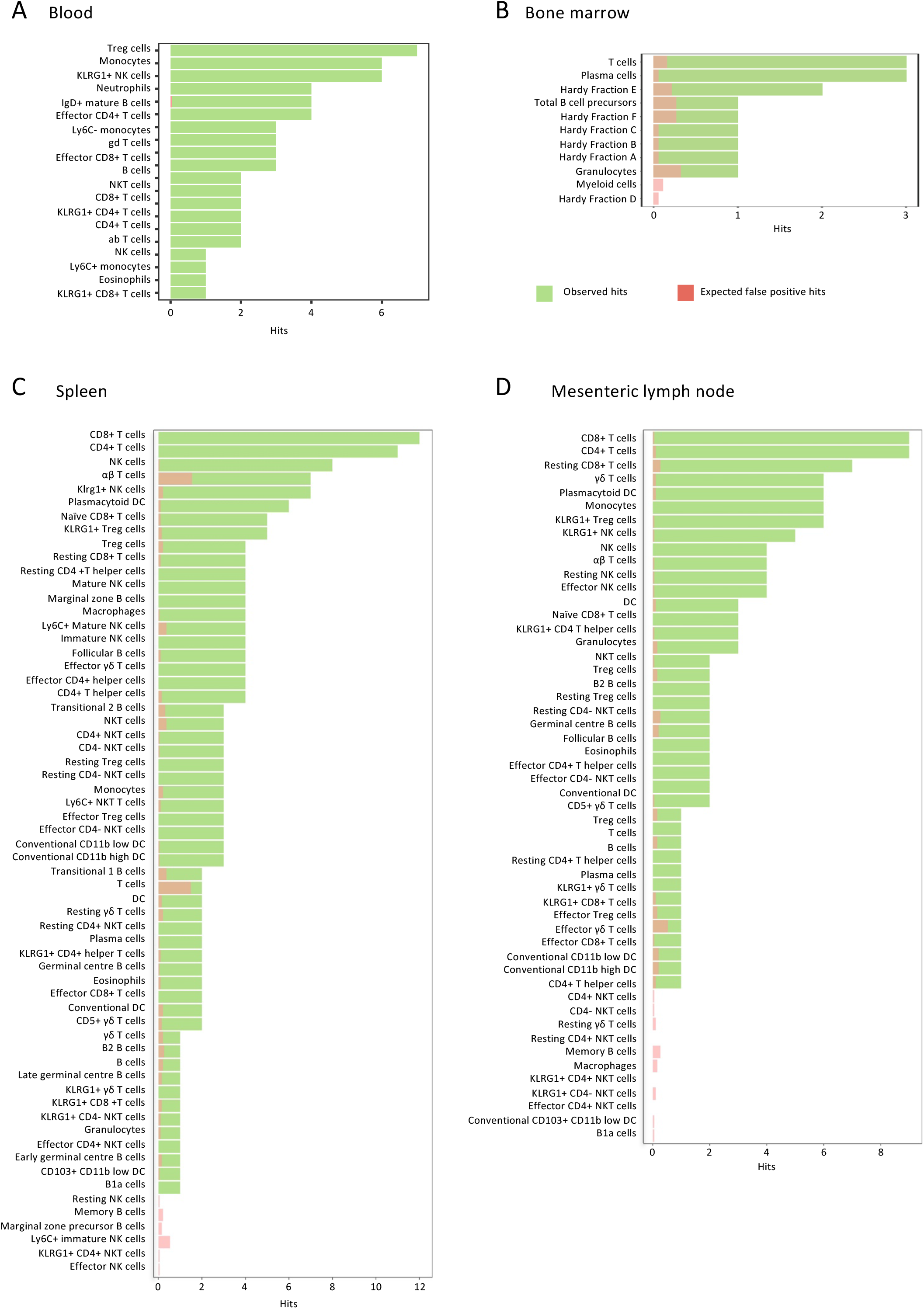
False positive rates are far below hit rates. A-D Expected number of false positives estimated by simulation (re-sampling of wt controls) compared to the number of significant phenotypes detected in: (A) PBL (n>450 per sex), (B) BM (n>350 per sex), (C) SPL (n>250 per sex), and (D) MLN (n>250 per sex). Observed hits depicted in green; expected false positives in red.

**Supplementary Figure 4:**
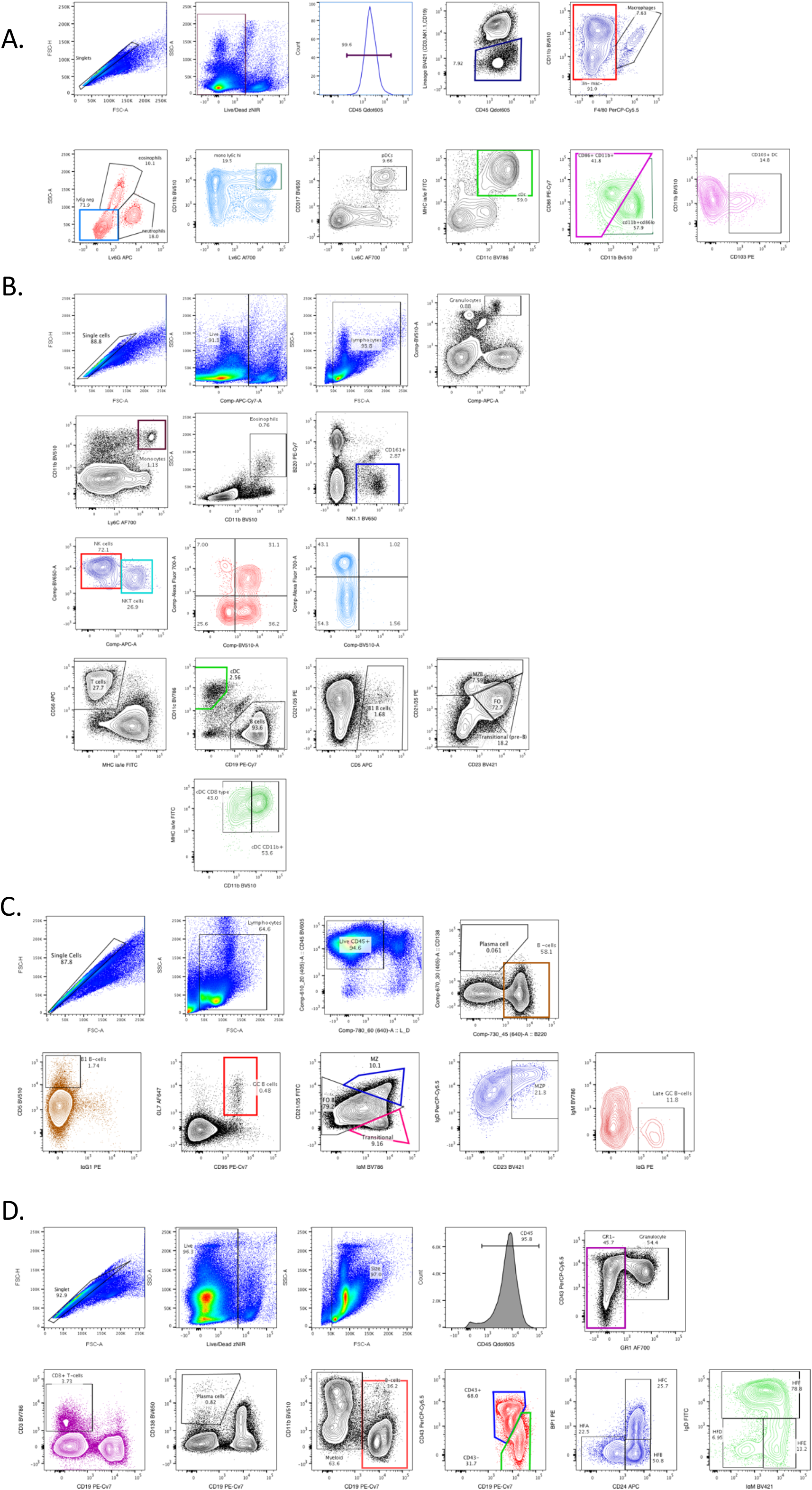
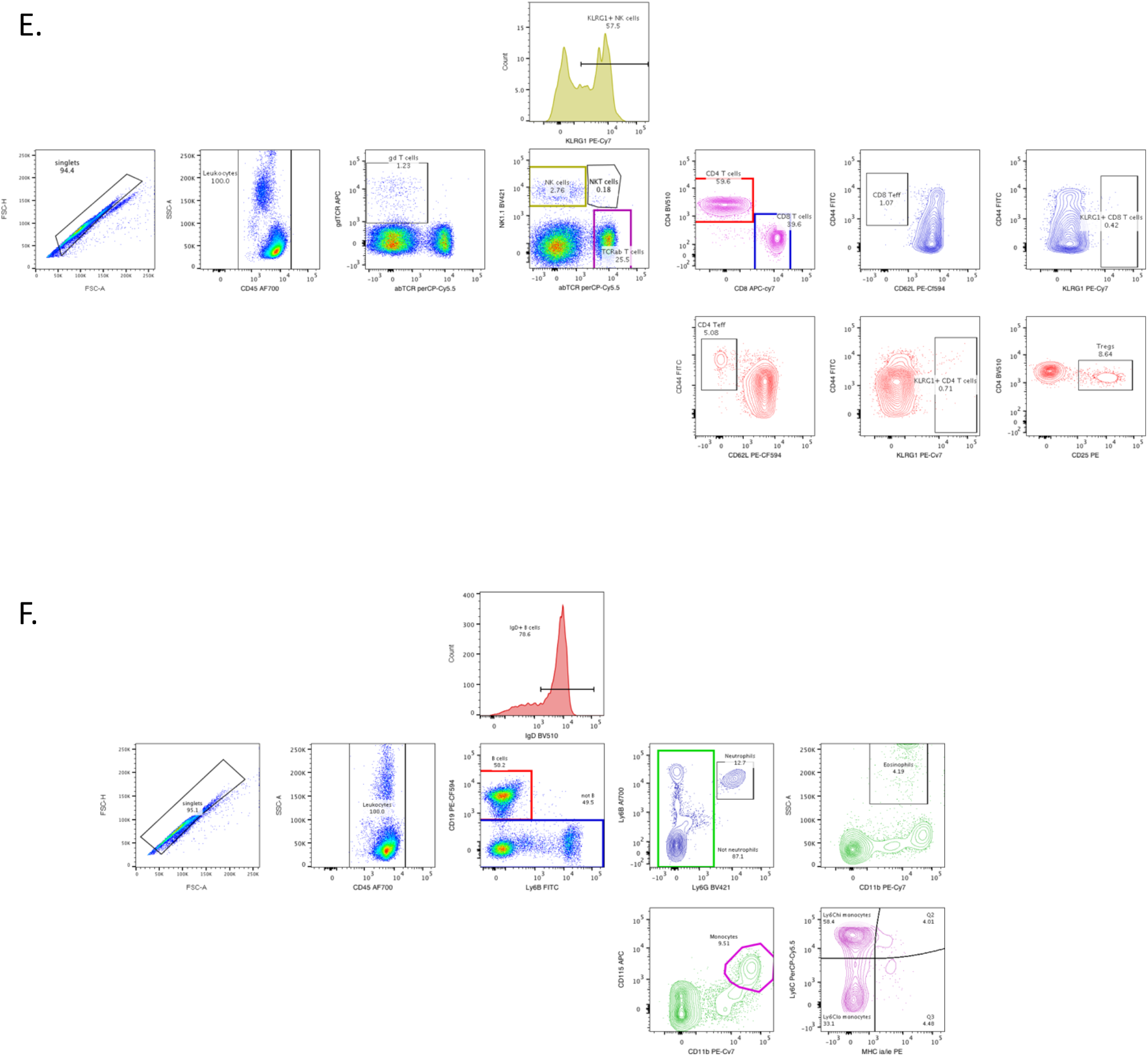
Flow cytometry gating strategies. Subsets of key populations are shown in gates of the same colour. Percentages denote percentages of parent population. Panels can be found in Table S1. Representative data for: A. Myeloid cells (data from spleen) B. B-cells NK cells (data from spleen) C. B-cells (data from spleen) D. Bone marrow E. T/NK cells (data from blood) F. B cell/monocyte (data from blood)

